# A Fast Interferometric Beam Shaper for Multi-Emitter 3D MINFLUX

**DOI:** 10.1101/2023.12.09.570565

**Authors:** Maximilian K. Geismann, Alba Gomez-Segalas, Alessandro Passera, Mehrta Shirzadian, Francisco Balzarotti

## Abstract

Beams of light that feature an intensity zero are essential to a variety of optical microscopy methods. Super-resolution techniques like STED and RESOLFT, together with localization strategies like MINFLUX and MINSTED, rely on accurate and fast displacements of such beams and their zeros. Extending these methods to the third dimension requires axial deflection, which, in contrast to lateral deflection, remains technologically challenging on the microsecond scale. Here, we present a fast general-purpose beam-shaping polarization interferometer that, instead of displacing the entire beam, enables such axial deflections by deforming the beam shape to deflect its zero. Based on this approach, we showcase a four-channel dual-color excitation system for three-dimensional MINFLUX imaging and tracking. We include first demonstrations of improved MINFLUX localization schemes that utilize the combination of distinct beam shapes and three-dimensional multi-emitter tracking. We believe that the presented approach will facilitate the broader adoption of three dimensional MINFLUX and provides a versatile basis for future implementations of advanced single-molecule localization methods.

## Introduction (without heading)

Displacing and deforming beams of light in a controlled manner is integral to a wide range of methods such as optical tweezers and microscopes. Super-resolution fluorescence imaging techniques, such as STimulated Emission Depletion (STED)^1^, and REversible Saturable Optical Linear Fluorescence Transitions (RESOLFT)^2,3^, as well as localization strategies like Maximally INFormative LUminescence eXcitation (MINFLUX)^4^ and MINSTED^5^, require displacing beams that feature zeros of intensity. These zeros—that either constrain regions from which fluorescence can be emitted or generate a spatially variant signature—enable the precise assignment of a fluorophore’s location with respect to a targeted coordinate.

MINFLUX^4^ stands out as a single-molecule localization strategy, distinguished by its approach of sequentially probing the position of an emitter with different beam shapes. Its high localization efficiency—the number of emitted photons required to reach a given localization precision—has been harvested for stochastic super resolution imaging^6,7^ and single-molecule tracking, reaching spatial and temporal resolutions in the nanometer^8^ and sub-millisecond^9–11^ scales. MINFLUX localizations in one-, two- and three-dimensions have been achieved through beams shaped by half-moon^10^, vortex^4^ and top-hat^8^ phase masks. Even Gaussian beams^4^ are efficient when used appropriately.

Irrespective of shape, the beams employed by the above methods require displacement. Available lateral scanning technologies include galvanometric, piezoelectric, electro-, and acousto-optic deflectors, while axial scanning technologies include electro-, acousto-optic, and liquid lenses. MINFLUX implementations have been based on electro-optical lateral deflection^4^, as it is compact, achromatic, simple to use, and delivers sub-nanometer and sub-microsecond deflection. For axial deflection, previous implementations resorted to an electro-optically tunable lens^8^or a deformable mirror^12^. These devices come at an increased cost and exhibit more complexity than their lateral-deflection counterparts. The tunable lens has a limited focal length of down to one meter and a response in the order of 30 µs, limited by its high capacitance and the current provided by the driving high-voltage amplifier. Deformable mirrors, being micro-electro-mechanical devices, face similar speed constraints.

We present an interferometry-based strategy for fast beam shape manipulation where the coherent superposition of specific pairs of beams with different shapes results in displacements or tilts of point-, line- and plane-like zeros of intensity, making it attractive for super-resolution techniques such as STED and RESOLFT, localization techniques like MINFLUX and MINSTED, and optical tweezer applications. To realize this concept we developed and characterized a general-purpose four-channel polarization interferometer compatible with continuous-wave and pulsed beams. It is the key component facilitating 3D and two-color operation for imaging and tracking in the presented MINFLUX system and we believe it will make these features more accessible for future MINFLUX implementations. Furthermore, this technology enabled the first demonstrations of MINFLUX excitation schemes mixing distinct beam shapes for improved precision and concurrent two-emitter MINFLUX tracking in 3D, following the recent introduction of simultaneous two-emitter MINFLUX tracking in 2D^13^. These advancements pave the way for multi-emitter MINFLUX tracking as a method for studying the kinetics and conformational states of flexible protein complexes free from range limitations present in single-molecule FRET^14^ and with time resolutions in the sub-millisecond range, surpassing high-speed AFM^15^.

## Results

### Interferometric manipulation of an intensity zero point, line and plane

Regions of space where focused beams have no intensity exist as points, lines or planes and can be generated by distinct phase masks at the back focal plane of a lens. Near these locations with no intensity, the spatial dependence of the electric field and its polarization take a variety of forms. It is possible to manipulate the shape and location of these regions by interferometrically combining distinct beams, thus mapping their relative amplitudes and phases to spatial features. Figure 1 displays various simulated and measured beam combinations that displace and tilt zero-intensity regions (see methods, PSF simulation and measurement).

**Fig. 1:**
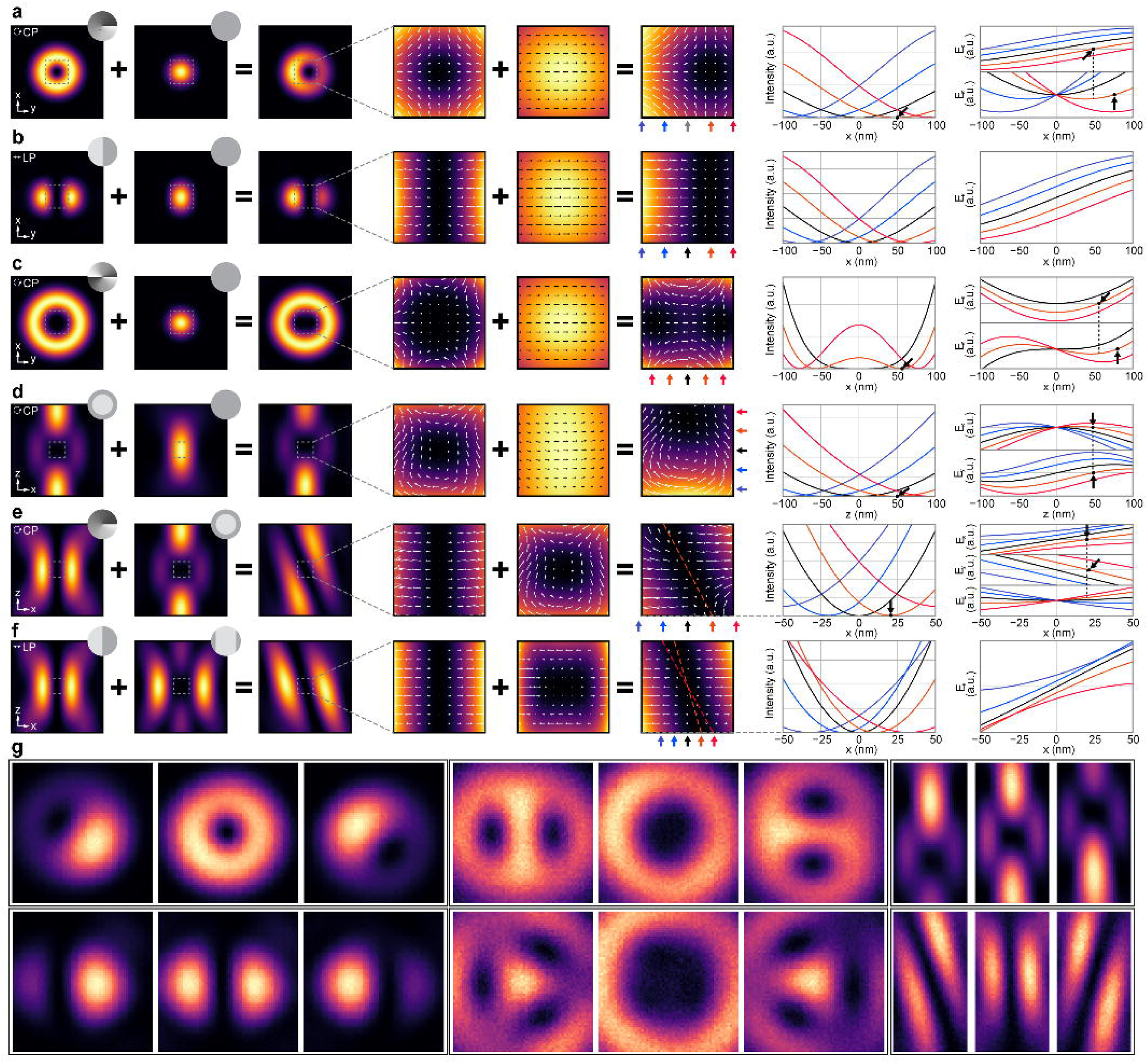
Interference of canonical beams. **a**, Combination of charge-1 donut with Gaussian beams. **b**, Combination of bi-lobed and Gaussian beams. **c**, Combination of charge-2 donut and Gaussian beams. **d**, Combination of bottle and Gaussian beams. **e**, Combination of charge-1 donut and bottle beams. **f**, Combination of bi-lobed and bottle-slice beams. (Left column) Intensities of combined beams at the xy (a-c, 800x800nm) and xz (d-f, 1200x1200nm) plane. CP: circularly polarized. LP: linearly polarized. Dashed squares: 200x200nm. The corresponding phase masks for each canonical beam shape are indicated in grey at the top right corner. (Middle column) Zoomed intensities in the dashed squares with a vector plot of the xy (a-c) or xz (d-f) components of the electric field. The resulting displaced zero coordinate is marked by an arrow. (Right column) Intensity and electric field profiles for each x, y and/or z component along the x axis (a-c), z axis (d) or x axes offset in z (e,f), chosen to highlight the interplay between electric field components. Mismatches between zeros of different components are marked with arrows. **g,** Interferometrically shaped PSFs measured on 40 nm gold particles via scattering for three different mixture amplitudes. Top left: donut plus Gaussian, xy; Top middle: charge-2 donut plus Gaussian, xy; Top right: Bottle beam plus Gaussian, xz; Bottom left: bi-lobed plus Gaussian, xy; Bottom middle: charge-3 donut plus Gaussian, xy; Bottom right: donut plus bottle beam, xz.

Several of these combinations are based on matching pairs of beams with linear and uniform electric fields, see Table 1. For example, a circularly polarized donut beam, created using a 2*π* vortex mask, features an axial zero line and a linearly dependent electric field across the xy plane that can be laterally displaced when combined with a circularly polarized Gaussian beam (Fig. 1a, Supplementary Fig. S1). As the donut beam contains all phases of electric field at any given time, there will always be a location where it destructively interferes with the Gaussian beam. That location is dictated by the relative amplitude and phase of the Gaussian beam, mapping them to physical polar coordinates^16^ (Supplementary Fig. S2). A similar phenomenon occurs with a linearly polarized beam shaped by a half-moon phase mask, creating an axial zero plane that shifts laterally when paired with a linearly polarized Gaussian beam (Fig. 1b, Supplementary Fig. S3). When combining a charge-2 donut beam with a Gaussian beam (Fig. 1c, Supplementary Fig. S4), the axial zero line at the origin splits into two zero lines, opposite from each other, which are also mapped into polar coordinates by the relative phase and amplitude of the beams (Supplementary Fig. S5).

**Table 1.** Canonical beam shapes with their polarization state, corresponding back-focal plane phase mask and zero type. Second order approximations of the focused electric field at the origin are shown for each component of the electric field in the last three columns, excluding proportional constants. The constant *c* absorbs differences between lateral and axial prefactors.

| Beam | Polarization | Phase mask | Zero type | $E_x(\vec{r})$ | $E_y(\vec{r})$ | $E_z(\vec{r})$ |
| --- | --- | --- | --- | --- | --- | --- |
| Gaussian | Circular | Flat | None | 1 | $i$ | $x - iy$ |
| Gaussian | Linear x | Flat | None | 1 | 0 | $x$ |
| Bi-lobed | Linear x | Half-moon | Transverse plane | $x$ | 0 | $-ixy$ |
| Donut charge 1 | Circular | Vortex $2\pi$ | Axial line | $x + iy$ | $ix - y$ | $-x^2 + y^2 + 2ixy$ |
| Donut charge 1 | Circular | Vortex $-2\pi$ | None | $x - iy$ | $ix + y$ | 1 |
| Donut charge 2 | Circular | Vortex $4\pi$ | Axial line | $-x^2 + y^2 + 2ixy$ | $i(-x^2 + y^2) - 2xy$ | 0 |
| Bottle | Circular | Top-hat | Point | $x^2 + y^2 + cz$ | $i[x^2 + y^2 + cz]$ | $ix - y$ |
| Bottle-slice | Linear x | Top-hat slice | Lateral line | $x^2 + cz$ | 0 | 0 |

Furthermore, interferometric combinations that specifically alter the axial distribution within the beams can be engineered. For instance, the central zero point of a bottle beam can be axially shifted by combining it with a Gaussian beam (Fig. 1d, Supplementary Fig. S6), as the bottle beam is purely laterally polarized across the z-axis with a linear dependence on z. Other innovative combinations include tilting the axial zero line of a circularly polarized donut beam (Fig. 1e, Supplementary Fig. S7,Supplementary Fig. S8) and tilting the zero plane of a linearly polarized bi-lobed beam (Fig. 1f, Supplementary Fig. S9) when combining them with the bottle beam and its linear analogue the bottle-slice beam, respectively. See Table 2 for a list of the beam combinations presented.

**Table 2.** Examples of some beam combinations and the resulting effects on the featured zero.

| Beam 1 | Beam 2 | Polarization state | Zero type | Interference Result |
| --- | --- | --- | --- | --- |
| Donut charge 1 | Gaussian | Circular | Axial line | Lateral displacement |
| Bi-lobed | Gaussian | Linear | Axial plane | Lateral displacement |
| Donut charge 2 | Gaussian | Circular | Axial line | Lateral displacement<br>Double lines |
| Bottle | Gaussian | Circular | Point | Axial displacement |
| Donut charge 1 | Bottle | Circular | Axial line | Line tilt |
| Bi-lobed | Bottle-slice | Linear | Axial plane | Plane tilt |

However, beams focused with high numerical aperture contain axial electric fields^17^, leading to a zero degradation if (i) the interferometric combination does not yield a null axial component or, if (ii) it does, but its location does not match the zero of the lateral components (see electric field profiles in Fig. 1a-g with zero mismatches marked by arrows). Additionally, a mismatch of Gouy phases, as in the case for the bottle beam and the Gaussian, degrades axially shifted zeros (Supplementary Fig. S10). Such degradation manifests in scenarios involving large displacements of the zero and depends on the numerical aperture of the system and fill factor of the beams.

Despite these circumstances, the interferometric shifting of zeros can imitate whole beam deflection for techniques, such as MINFLUX, where only the parabolic intensity distribution around a zero matters. It offers an enticing alternative to current technologies for implementing axial deflection as discussed in the following section.

### A four-channel common-path beam-shaping polarization interferometer

We present an implementation of our interferometric beam shaping strategy with a system that is compact, modular, scalable, cost-efficient, flexible, and capable of dynamic manipulation. The system comprises three main blocks (Fig. 2a-c): a polarization rotator, a beam shaper, and a beam combiner that control the interference of two distinct beam shapes by modulating their relative amplitudes. In this two-color design, these components act on four independent paths, two per laser wavelength (Fig. 2d).

**Fig. 2:**
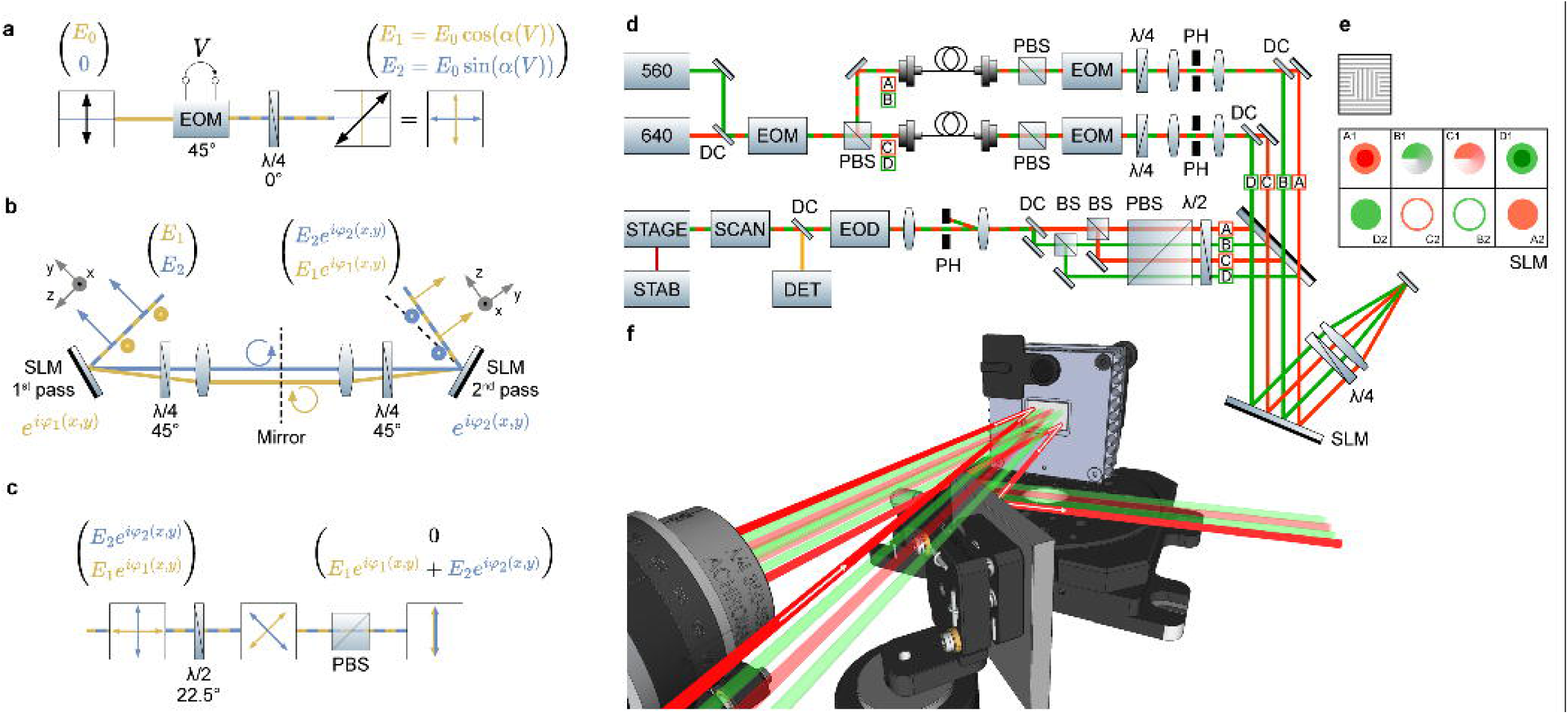
Implementation of a multi-channel polarization interferometer. **a**, Polarization rotation module. The square with arrows represents the polarization state. **b**, Simplified diagram of the beam shaping module. A double-pass configuration on an SLM is shown unfolded with respect to its mirror. **c**, Beam combination/interference module. **d**, Complete system. **e**, Detailed grating structure used in all SLM regions (above), example phase masks (below) for interferometric bottle beam deflection in A1+A2 (and D1+D2) and donut beam generation on B1 (and C1) for red (and green) wavelengths; B2 and C2 are unused. **f**, 3D render of the SLM with all beam paths. The arrows indicate the propagation direction for one of the red paths (path A): it passes above a two-inch square mirror glued on a half-inch kinematic mount, reflects off the SLM, propagates through the imaging system and the QWP (not shown), reflects off the SLM again at a different height and is eventually picked up by the square mirror.

The first block, a polarization rotation module (Fig. 2a), allows us to change the relative amplitudes of the beams that will interfere. The rotator is comprised of an Electro-Optical Modulator (EOM) with its principal axis at 45° and an achromatic Quarter-Wave Plate (QWP) at 0°. For a linearly polarized Gaussian beam 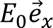 at the input, the output is a rotated linearly polarized state 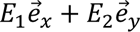 (shown in yellow and blue) with *E*_1_ = *E*_0_ cos *α*(*V*) and *E*_2_ = *E*_0_ sin *α*(*V*), where the rotation angle depends on *V*, the voltage applied to the EOM. This combination offers maximal modulation speed and has no chromatic angular dispersion. Implementing fast phase modulation, e.g., to realize the zero rotations depicted in Supplementary Fig. S2, Supplementary Fig. S5 and Supplementary Fig. S8, only requires an additional EOM with a 0° principal axis.

A double-pass configuration on a Spatial-Light Modulator (SLM) independently shapes the wavefront of each orthogonal linear polarization state (in yellow and blue), depicted in fig. 2b as a split diagram around a metallic mirror. The SLM first imprints a programmable retardation *φ*_1_(*x*, *y*) on *E*_1_ (yellow beam), while the light in the orthogonal polarization state (blue beam) remains ideally unaffected. The following elements—QWP, convergent lens and metallic mirror—produce an inverted image of the SLM onto itself, as the system behaves as a 4*f* imaging system, and a 90° rotation of the polarization. This makes the second pass fall on a distinct region of the SLM (not shown in the simplified diagram of Fig. 2b), adding an arbitrary retardation *φ*_2_(*x*, *y*) to *E*_2_ (yellow beam) and resulting in an output 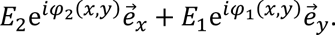

Finally, both patterned fields *E*_1_and *E*_2_undergo a 45° polarization rotation by a half-wave plate and a Polarizing Beam Splitter (PBS) produces the final interference (Fig. 2c), outputting 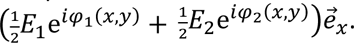 Both interfering beams share their optical path, meaning that this system has no need of active feedback loops or adjustments to maintain the relative phase stability.

It is common practice to use a blazed diffraction grating (e.g., horizontal) on SLM holograms to reject undesirable spurious phase structures imprinted on its zero-th order, i.e., the direct reflection^18^. A common spatial filter for the four paths (shown in Fig. 2d) rejects the zero-th order while the first diffraction order, which carries the patterned beams, passes through. As an added benefit, the use of the first diffraction order in the double-pass configuration filters out the crosstalk between polarization states due to, for example, the imperfect retardation of the QWP, see Supplementary Note 1. Additionally, we apply an orthogonal grating outside of the selected pupil region (Fig. 2e) for suppressing spurious light.

The modular geometry described above is replicated to produce four independent paths, each carrying an interferometric mixture of two shaped beams on a single SLM (Fig. 2d), with only one path active at any given moment. We tailored this system to operate with two paths in the green range (560 nm) and two in the red range (640 nm). After combining both laser sources with a dichroic mirror, an EOM-based path switching selects paths A/B or C/D for red/green. Each path contains its own EOM-based polarization rotator (as in Fig. 2a) followed by a spatial filter for cleaning up the mode. Paths A/B and C/D are split by dichroic mirrors and reach distinct regions of the SLM labeled as A1/B1/C1/D1 (Fig. 2e). The double-pass configuration changes their height and inverts the order of the regions on the second pass to D2/C2/B2/A2. Ultimately, the system can realize any of the beam manipulations previously mentioned in any of the four paths. After interference at the subsequent PBS, all paths are recombined into one and propagate into a MINFLUX confocal microscope, comprised of the commonly required blocks of scanning, detection, and stabilization (see Methods, MINFLUX Microscope).

In conclusion, for each excitation wavelength it is possible to select one of four phase masks e.g., *φ*_*A*1_, *φ*_*C*1_, *φ*_*A*2_, *φ*_*C*2_ and, combine two pairs of them interferometrically e.g., *E*_0_[*χ*_*A*_e^*iφ*_*A*1_^ + (1 − *χ*_*A*_)e^*iφ*_*A*2_^] and *E*_0_[*χ*_*C*_e^*iφ*_*C*1_^ + (1 − *χ*_*C*_)e^*iφ*_*C*2_^], where the mixture coefficients *χ*_*A*_ and *χ*_*C*_ are dictated by the polarization rotation. We exploit this flexibility in the next section.

### A fast axial scanner for efficient 3D MINFLUX

MINFLUX localization requires sequentially probing the emission of a fluorophore with tailored excitation beam shapes at distinct locations. These MINFLUX configurations usually enclose a circular or spherical region with a diameter *L* that scales the achievable localization precision and is shrunk sequentially in iterative MINFLUX^8^. Our system can produce a variety of such configurations, some of which are shown in Fig. 3a: all Gaussian or donut beams for 2D localization, all deformable bottle beams for 3D localization and mixtures of donut and bottle beams for improved photon efficiency, as well as interleaved wavelengths for simultaneous MINFLUX localization in distinct spectral bands (Fig. 3b). While the interferometric approach can produce lateral deflections suitable for MINFLUX (Figure 1a,g), we employed a pair of electro-optical deflectors that surpasses its range at a similar cost with a simpler implementation. To achieve axial deflection (Fig. 1d), however, this approach stands out for its accessibility and unrivaled time response.

**Fig. 3:**
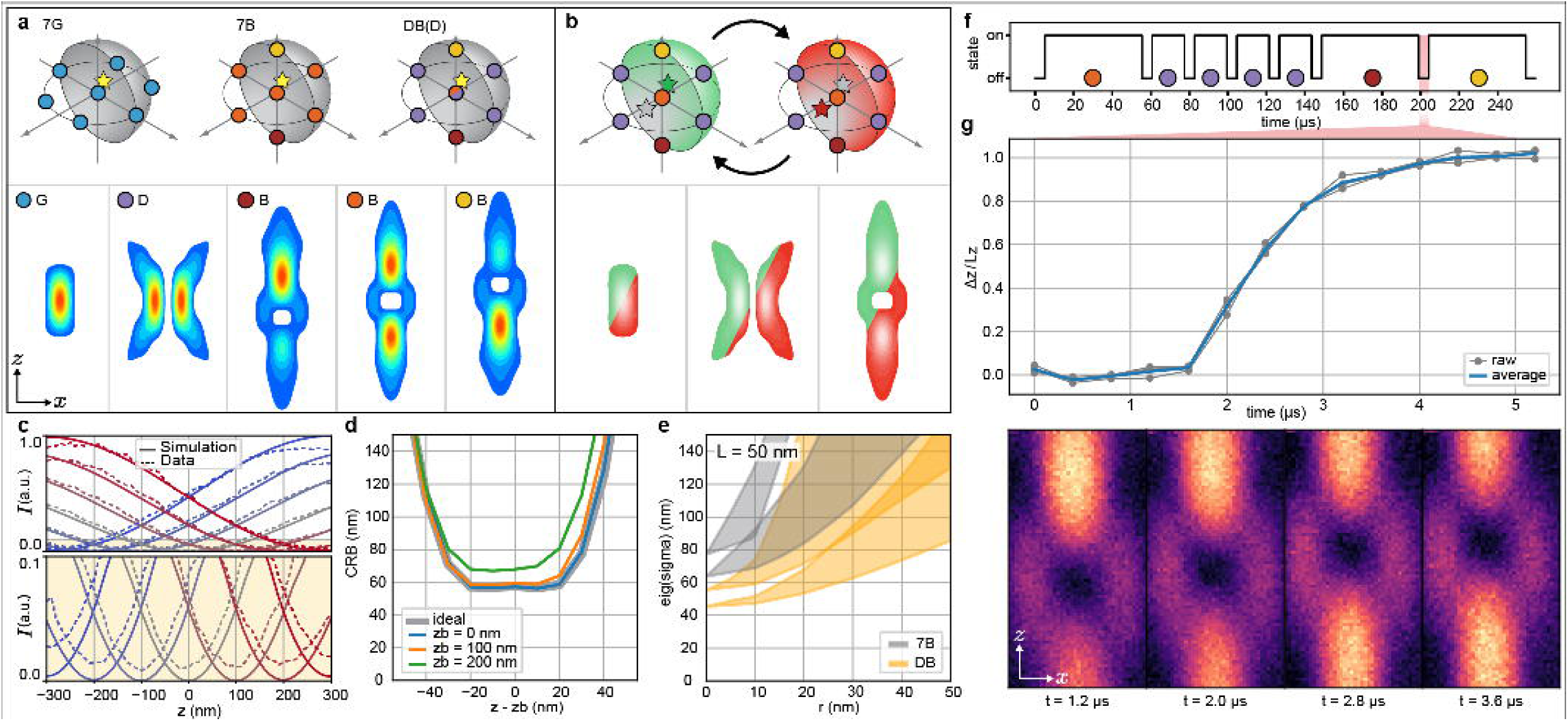
MINFLUX excitation schemes with interferometrically deformed beams and mixed shapes. **a**, Depiction of the 2D and 3D MINFLUX excitation schemes realized with our system (top) and their corresponding beam shapes (bottom): Gaussian beam (G; blue), donut beam (D; purple) and bottle beam (B; red, orange, yellow) at different interferometric deflection states. **b**, Strategy for tracking two emitters simultaneously by rapidly switching between schemes for red and green excitation wavelengths, with seven exposures each. **c**, Juxtaposition of simulated and measured intensity profiles of the deflected bottle beam zero. Bottom: zoom-in of the yellow area. Despite the deformation, the profiles stay parabolic over a large range. The zero degradation of the simulated profiles is visible towards the edges. The measured profiles never reach a true zero due to background and aberrations. **d**, Comparison of the CRB for a single photon at a signal-to-background ratio of five along the optical axis z for ideally deflected bottle beams (without deformation, gray) and for our interferometric approach. Only when localizing emitters far off focus, given by the axial distance zb, the performance degrades slightly. **e**, Eigenvalue ranges of the 3x3 CRB covariance matrix for one collected photon with the *mixed beam* scheme comprised of donut and bottle beams (DB) versus the conventional 3D MINFLUX excitation scheme consisting of bottle beams only (7B). Each condition displays two shaded areas representing the range of the minimum (lower) and the maximum (upper) CRB eigenvalue for emitter positions on a sphere with radius r centered on the excitation pattern, indicating anisotropy; the width of each area accounts for spatial inhomogeneity. **f**, Timing diagram for the mixed beam scheme, with larger weights for the bottle beam exposures. **g**, Measurement of the interferometric deflection transient for a distance of 271 nm (top). Each point corresponds to the location of the zero of intensity obtained from a complete PSF measurement (bottom, selected time points).

Interferometric axial deflection of the *zero point* of a bottle beam overlapped with a Gaussian beam is shown via single-molecule fluorescence Point Spread Function (PSF) measurements (see Methods: PSF simulation and measurement) alongside simulations in Fig. 3c. This zero level can degrade due to a lack of coherence (see Supplementary Note 2) and misalignment between the interfering beams (Methods: Four channel polarization interferometer setup). Even under ideal conditions, a mismatch of the Gouy phase of both beams slightly degrades the deflected zero’s quality as discussed in the previous section. Remarkably, the interferometer also works for pulsed excitation lasers as shown in Extended Fig. 1.

MINFLUX excitation schemes that displace beams with a zero of intensity harvest a gradient of *photon collection probability* that leads to a high localization precision^19^. This is hampered by zero degradation and is equivalent to a loss of signal-to-background ratio. Fig. 3d features a comparison of localization Cramér-Rao Bound (CRB) between defocused bottle beams with perfect zeros and interferometrically deformed bottle beams with zeros that degrade depending on their position. Despite the imperfect zeros there is no major deviation from the ideal localization precision.

Displaced bottle beams provide axial information, with poorer performance than donut beams in the lateral direction. This is because the emitted background *per emitted fluorescence photon* is higher for bottle beams than for donut beams, due to their weaker curvatures, larger size, and axial lobes. One could consider working in low background scenarios, but MINFLUX operates by purposely lowering the signal in exchange for high information content per photon by shrinking *L*. Thus, one should always strive to work in a regime in which background and/or zero quality are not negligible. We therefore implemented the capability of using a mixture of donut and bottle beams within one MINFLUX excitation scheme for lateral localization and axial localization, respectively. A CRB comparison of both scenarios (Fig. 3e) reveals a significant improvement for a molecule localized throughout the volume of the excitation pattern for an *L* of 50 nm.

The speed of the interferometric deflection dictates the wait time between the axial exposures of a MINFLUX excitation scheme (Fig. 3f) and it is limited by the bandwidth and electrical current of the amplifier driving the EOM of the polarization rotator (Fig 2a). We determined the rise time of the axial deflection by imaging the transient PSF at a time resolution of 200 ns (Fig. 3g, Methods: Bottle beam deflection speed Characterization). We measured a rise time (10%-90%) of <1.5 µs and a stabilization time (<1% of the asymptote) of <2.5 µs, with a system delay of 1.6 µs.

### MINFLUX imaging and tracking

For proof-of-principle demonstrations of the system we used static and dynamic DNA origami structures as well as fixed cell samples. First, we designed and folded a DNA origami that arranges 12 Alexa Fluor 647 dye molecules in a three-by-four grid with 20 nm spacings for 3D STORM imaging (see Methods). The results are shown in Fig. 4a: all sites are clearly resolved with a Full Width at Half Maximum (FWHM) of the fitted Gaussian distribution of <3 nm in xy and <4.5 nm in z corresponding to standard deviations for the localization precision of <1.3 nm and <1.9 nm, respectively. Averaging all emission events per site yields very high localization precisions as reported for DNA-PAINT^20^. On the STORM-MINFLUX data shown here, the FWHM of the average location ranges from 0.5nm to 1.0nm, corresponding to standard deviations between 0.2nm and 0.4nm in all dimensions. Furthermore, we acquired 3D images of the Nuclear Pore Complex (NPC) by labelling SNAP-tagged NUP96 with Alexa Fluor 647 in fixed U-2 OS cells, see Fig. 4b.

**Fig. 4:**
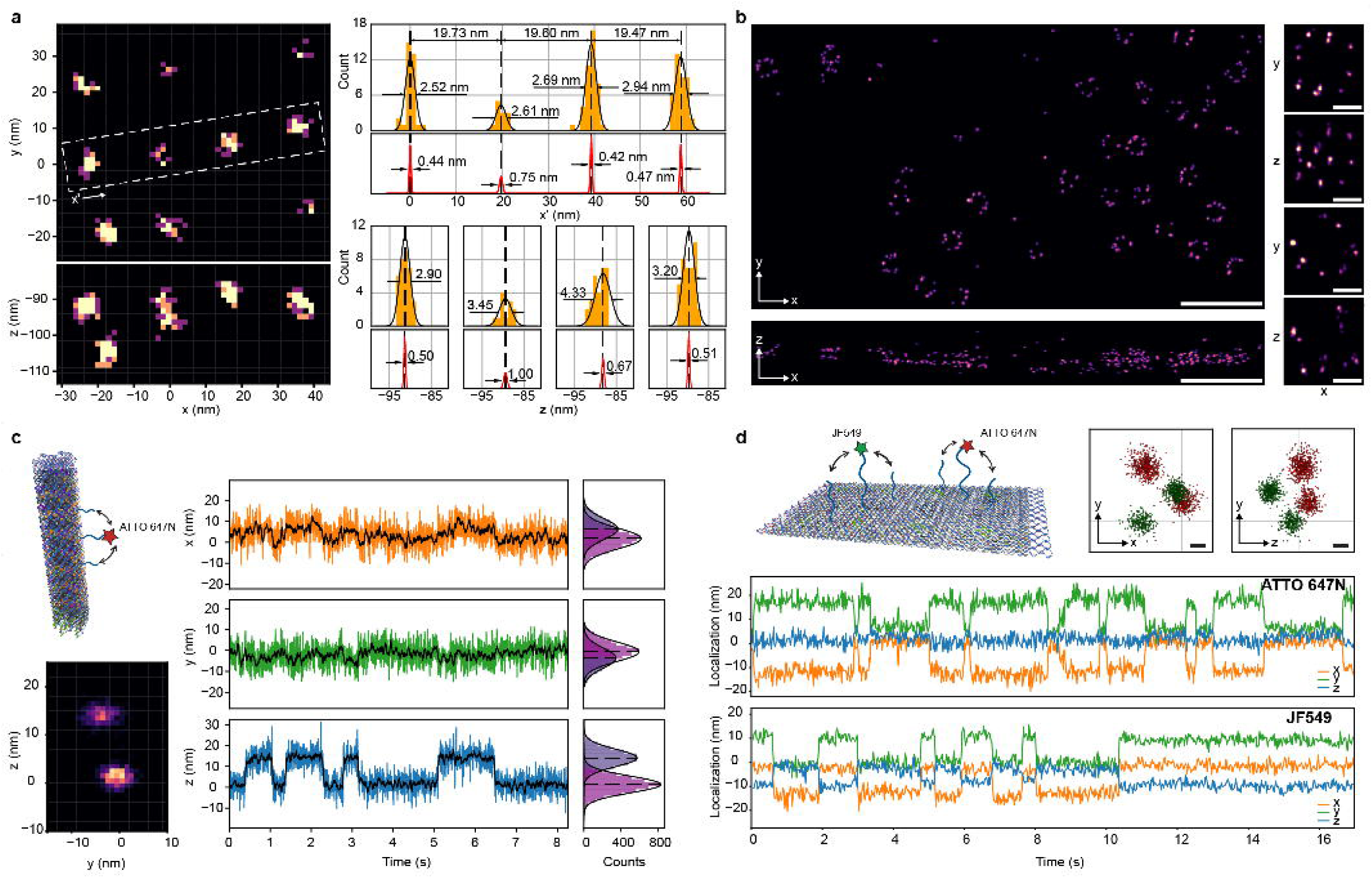
MINFLUX imaging and tracking. **a**, 3D Imaging of Alexa Fluor 647 fluorophores arranged in a 3x4 grid with 20 nm spacing on a DNA origami (left). Localization histogram along the marked region with indications for distances and FWHM (top right) and corresponding histograms along z for each site (bottom right); for each site, the FWHM of the mean of all ocurrences is marked in red. **b,** 3D imaging of NUP96-SNAP labelled with BG-Alexa Fluor 647 in fixed U-2 OS cells. **c**, 3D tracking of ATTO647N conjugated to a DNA strand that oscillates axially between two binding sites on a DNA origami (top left). Histogram of localizations at 10 ms temporal resolution (bottom left). Localization traces for each coordinate (middle) at 1 ms temporal resolution (x: orange, y: green, z: blue) and with 10-fold temporal binning (black). The origin of the axes is set by the excitation pattern center after the initial centering step. Histograms for the two distinct positions of the molecule (right). **d**, Schematic of a 2D origami (upper left) with ATTO647N and Janelia Fluor 549 dyes, each transiently binding to two distinct sites. XY and YZ projections (upper right) of all localizations in the traces displayed below. Localization traces for ATTO647N and Janelia Fluor 549 (bottom), at 20 ms and 25 ms temporal resolution, respectively. DNA origami renders were generated with oxView. Scale bars: (**b**) 500 nm for large field of view, 50 nm for single pores, (**d**) 5 nm.

To showcase the 3D tracking capability and temporal resolution of the system, we designed a DNA origami with a fluorescently labelled DNA strand oscillating axially between two catching sites 14 nm apart. The DNA origami itself is rod-like, approximately 50 nm-tall, and features 16 biotin-modified DNA staple strands for it to stand vertically on streptavidin-functionalized surfaces (Fig. 4c). Localization traces shown in Fig. 4c are at 1 ms and 10 ms temporal binning, which correspond to 150 and 1500 average photon counts per localization, respectively. At 1 ms temporal resolution, the standard deviations for the localization precisions are 4.9, 4.7 and 4.1 nm in x, y and z. After further binning, the measured distance between catching sites is 14.9 ± 1.7 nm, see Methods: Data analysis.

The measurement scheme for tracking consisted of a first step where the excitation pattern is actively centered onto the fluorophore via a proportional-integral closed-loop control system that uses the live estimated MINFLUX localization. This feedback loop can flexibly act on the tip-tilt mirror, the EODs, and the interferometric axial scanner simultaneously. We chose the former because it is de-scanned and hence the collection efficiency is optimal after centering. Once centered, the excitation pattern is static over time and probes the molecule until photobleaching. The effectiveness of the centering step is evident in the localization traces in Fig. 4c, where MINFLUX tracking is initiated within a 10 nm range from the molecule’s actual location.

Finally, we utilized the two-color excitation of the presented MINFLUX implementation to simultaneously track two spectrally distinct emitters (ATTO 647N and Janelia Fluor 549 dyes), each transiently binding to two distinct sites on a 2D DNA origami structure of programmable dynamics (Fig. 4d). The average photon count per localization in the displayed localization traces is 1200 for both emitters, resulting in 20 ms temporal resolution for ATTO 647N and 25 ms for Janelia Fluor 549. The corresponding standard deviations for the xyz-localization precisions are 2.2, 2.3, 1.8 nm for ATTO 647N and 1.8, 1.7, 1.6 nm for Janelia Fluor 549.

In this case, the measurement scheme involved a centering step at 640 nm excitation wavelength, converging onto the far-red dye, and defining a volume that encompasses both emitters. Following centering, the two excitation lasers were interleaved in time to probe the emitters with static excitation patterns, each one lasting approximately 250 μs. Notably, the sequence length is not limited by the deflection speed of either the EODs or the axial scanner presented here, but rather by the fluorophore’s emission rate and the need to bin sufficient photon counts to achieve single-digit localization precision.

## Discussion

We presented an interferometric strategy to manipulate a variety of focused beams, including some that are newly described. Our efforts centered on beams with intensity zeros and the manipulation of these regions through displacements and tilts. We proposed the use of three general optical blocks to realize this strategy: a polarization rotator, beam shaper and interferometer. An implementation of these blocks in a four-channel two-color configuration was introduced and employed as the excitation module of a custom MINFLUX microscope. We utilized this microscope for super-resolution imaging as well as single- and multi-molecule tracking with novel MINFLUX excitation schemes and dual-color operation, reaching state-of-the-art localization precision and time resolution.

The proposed common-path polarization interferometer is notable for its phase stability without active feedback, its versatility and extendibility. Its implementation is devoid of mechanical parts and manipulates beams at the microsecond time scale even for pulsed lasers. Nevertheless, slower and more economic polarization rotation technologies are also compatible, such as mechanical- or liquid-crystal-based methods. Simplified implementations could achieve similar interferometric effects. For example, a (commercially available) birefringent phase mask could replace the SLM. However, it is limited to a fixed wavelength and lacks the ability to correct aberrations or adjust phases. A single-pass SLM configuration could realize all interferometric combinations involving a Gaussian beam originating from the direct reflection of the modulator’s unaffected polarization. Nevertheless, this approach excludes blazed gratings, which would eliminate zero-order effects arising from the SLM pixelation^18^. Additionally, the lack of fine alignment and aberration correction on the directly reflected Gaussian beam leads to zero degradation, non-linear axial calibration and unintended tilts.

A wide range of methods might benefit from the presented approach. For instance, the volumetric neuronal imaging technique presented in^21^: instead of alternating two axial positions of the employed Bessel-droplet excitation foci via an SLM update, the interferometer could scan them continuously on a much faster time scale—potentially reaching the nanosecond regime when using a resonant EOM— to yield a more homogeneous distribution (see Supplementary Fig. S12). The same concept applies to optical trapping techniques that confine multiple particles axially^22^, and the axial deformation of the bottle beam might be used as a photophoretic trap capable of loading and unloading a particle similar to^23^. Furthermore, applying our axial displacement strategy to a STED beam is optimally suited for extending conventional fast STED imaging^24^ (especially for small volumes) and MINSTED^5^ into the third dimension.

We believe that this work will prove especially valuable for future MINFLUX systems: among MINFLUX’ major strengths is its isotropic localization precision in all three spatial dimensions. Yet, custom systems refrained from 3D implementations, presumably due to their complexity. Our method is not only a convenient alternative to current technologies for axial scanning but also unrivaled in its deflection speed. It integrates and scales well with other modalities, allowing the use of multiple beam shapes, several wavelengths and pulsed operation compatible with pulsed interleaved MINFLUX^25^. As such, we project that it will facilitate broader adoption of 3D MINFLUX and help to extend its current capabilities even further, unlocking new possibilities for investigating complex dynamic processes and structural studies on the molecular level involving single or multiple emitters.

## Acknowledgements

We thank Karina Arellano Ayala for assistance with sample preparation. We thank Anna Deputatova for assistance with designing and folding the 32HB origami. F.B. was supported by Boehringer Ingelheim. This project has received funding from the European Research Council (ERC) under the European Union’s Horizon 2020 research and innovation program (Grant agreement No. 853348). For the purpose of Open Access, the authors have applied a CC BY public copyright license to any Author Accepted Manuscript (AAM) version arising from this submission.

## Author Contributions

M.K.G., A.G. and F.B. designed and built the optical system, and programmed the hardware control. M.K.G., A.G., A.P., M.S. and F.B. programmed the processing and visualization software. M.K.G. and A.G. performed experiments and data analysis. A.P. designed and folded DNA origami and optimized used protocols. A.P. prepared and optimized cell samples. M.S. performed simulations and optimizations. M.K.G., A.G., A.P. and F.B. wrote the manuscript. F.B. supervised the project.

## Methods

### PSF simulation and measurement

Electric field calculations near the focus of an objective with high numerical aperture as shown in Fig. 1**a**-**f** and Fig. 3**a**-**c** were performed via a custom python implementation of ^26^.

The PSFs of the excitation beams were measured via a “bead stack” by moving the beam in xy via either the EODs or the tip/tilt mirror and in z by moving the fine stage (see Supplementary). The “bead” took the form of (i) fluorescent beads (see Fluorescent microspheres samples), (ii) single dye molecules attached to a DNA origami (see Origami sample preparation) or (iii) gold particles where the signal was the scattered excitation light (see Gold nanoparticles samples) detected on the avalanche photo diodes after removing the notch filter blocking the excitations wavelengths in the detection module.

The measured PSF gallery shown in Fig. 1**g** was acquired via scattering on gold particles. For the bi-lobed PSF the last QWP was removed to obtain linear polarization.

The measured intensity profiles shown in Fig. 3**c** were acquired on single molecule by scanning the voltage of the polarization rotator as well as moving the fine z-stage. One raw measurement contained 15 (not all are shown) profiles of different axial deflection offsets. Three such measurements were averaged and normalized to the maximum across all averaged profiles to produce the curves shown.

For the PSF measurements shown in Fig. 3**g** see Bottle beam deflection speed Characterization.

### MINFLUX Microscope

We use an EOM and a PBS to switch the main excitation beam carrying light of either 560 nm or 640 nm between two paths, each coupled into a fiber. The fiber outputs then go through a PBS for polarization cleanup and an EOM plus waveplates for polarization rotation. A three-lens (zoom) telescope with a spatial filter cleans the mode and allows for adjustable magnification. The two paths are then split by a dichroic mirror (DC) into four; two for 640 nm and two for 560 nm. All paths are aligned to be parallel and then go through the SLM double-pass configuration as described in the main text. All four paths are combined into one via regular beamsplitters and another DC. A telescope with an adjustable iris at its focal plane blocks the zero-th diffraction order and relays the image of the SLM. The remainder of the system adheres to the configuration presented in ^8^. For a detailed schematic see Supplementary Note.

The microscope control system comprises an FPGA board (NI USB-7856R), two data acquisition boards (NI PCIe-6363 and MCC USB-3114), and two Windows 10 PCs (one solely dedicated to the active stabilization). These hardware components are integrated through custom LabVIEW software, following the structure outlined in ^8^. However, we updated the software to specifically control the developed polarization interferometer and enable convergence onto molecules through the implementation of a proportional-integral controller in MINFLUX tracking experiments.

Furthermore, a Raspberry Pi (RPI 4B 8GB) running Raspbian is used to display phase masks on the SLM via HDMI. It runs a simple web server (written in Go) that receives the desired phase mask as a png file through a POST request which it then displays via a lightweight image viewer called “feh”.

### SLM setup

We used the phase linearization provided by the vendor but adjusted the scale for the wavelengths we employed: We calibrated the phase value that corresponds to a retardation of 2π (one full wavelength) by maximizing the power in the first diffraction order when scanning the amplitude of the blazed grating, i.e., the maximum phase of its periodic ramp.

To correct for excitation beam aberrations we employed a pupil-segmentation approach^27^ to measure the aberrated wavefront and apply resulting correction as an additive phase mask on the SLM. These corrections are crucial for achieving acceptable beam shapes, especially for the bottle beam.

The SLM was divided into 8 independent regions, each with its own calibration and correction.

### Four channel polarization interferometer setup

#### Polarization rotator

The output of all EOMs and their QWPs were monitored via a polarimeter (PAX1000VIS/M, Thorlabs) while continuously modulating at low frequency with a range larger than the lambda-half voltage. The rotation of the EOM was adjusted until the two linear polarization states occurring during the modulation were 90 degrees apart (180 degrees on the Poincaré Sphere) while the rotation of the QWP was adjusted to yield close to purely linear polarization states throughout the modulation.

#### SLM double pass

The SLM double pass was aligned such that the images of the first and the second pass coincide by applying the right pixel offsets to the SLM regions (according to the geometry) and adjusting the mirror (DP.MM). The rotation of the quarter-wave plate (DP.QWP) was adjusted to minimize crosstalk from one pass to the other. The rotation of the half-wave plate (DP.HWP) was set to 22.5° to have equal amplitudes in the mixture of polarization states leading to a linear deflection calibration, see Supplementary Fig. S14.

The voltage offsets on the path switch EOM (E.EOM) were adjusted to minimize crosstalk between paths of the same wavelength. The voltage values that select only the first or second pass were adjusted to minimize the crosstalk between the first and the second pass.

To tune the polarization interferometer of a path, we adjusted the relative (flat) phase between the first and the second pass, and the tip, tilt and defocus Zernike terms of the first pass on the SLM. For that, we set the voltage of the polarization rotating EOM to a value where both polarization components carry equal power measured on a power meter close to the objective. Then, plain gratings (aberration corrected) were displayed on the SLM, and the power was minimized (destructive interference) by adjusting the above terms. This procedure was automated.

For axial deflection of a bottle beam, a flat phase of π/2 was applied to the second pass as is necessary for the right phase relations between the top-hat phase mask and the Gaussian’s phase mask (plain grating). Finally, the quality of the deflection was tested by measuring intensity profiles along z on 100 nm fluorescent beads (FluoSpheres, 0.1 µm, Tetraspeck; Thermo Fisher Scientific) or single molecules on DNA origami (ATTO647N, Cy3B) and checking that the intensity minimum, the zero, was at an approximately constant level over a range of 600 nm for several deflection offsets. If not, small adjustments to the diameter of the π-phase step of the top-hat phase mask were made. Finally, the z-profiles were used to calibrate the polarization rotator’s EOM input voltage, such that, for a given target, the deflected distance matched the movement in z of the fine stage.

#### Beam combination alignment

All four paths were manually aligned by overlapping cross patterns generated on the SLM by simultaneously looking at an image- and a conjugate plane of the SLM. Fine corrections were made by adjusting the tip and tilt Zernike terms of the respective paths on the SLM while measuring intensity profiles along x, y and z directions on 40 nm fluorescent beads (FluoSpheres, 0.04 µm, dark-red fluorescence; Thermo Fisher Scientific) such that the intensity minima coincided.

### Bottle beam deflection speed Characterization

The xz-PSF measurement of the deflected bottle beam for several time points was achieved by collecting excitation beam photons scattered off 40 nm gold particles (see Gold nanoparticles samples) in a refractive index matched medium to avoid PSF artefacts caused by the reflection on the coverslip interface. An excitation scheme with two deflected bottle beams some axial distance apart (for Fig. 3: 271 nm) was used. We used the 3 µm diameter confocal detector path and set the excitation power to yield a count rate of roughly 8 MHz. The counts for each of the two excitations were collected in two separate channels. Then, for the second exposure the counts were gated with a 200 ns window that was shifted in time in steps of 400 ns to probe the transition from the axial position of exposure one to the axial position of exposure two. The z-trajectory of the gated channel was calculated by fitting the intensity minimum. Furthermore, it was drift corrected by fitting the intensity minimum of the ungated channel (static bottle beam).

### Sample preparation

#### Reagents

M13mp18-derived p7249 scaffold was obtained from NEB (catalog no. N4040S). Unmodified and biotinylated oligonucleotides for the rectangle origamis were obtained from MWG Eurofins, while unmodified and biotinylated oligonucleotides for the 3D origami and modified oligonucleotide for conjugation were obtained from Sigma-Aldrich. Freeze ‘N Squeeze columns (catalog no. 732-6165) were obtained from Bio-Rad. Cy3b-maleimide (catalog no. 11525834) was obtained from Cytiva. Alexa Flour 647-NHS ester (catalog no. 16820) was obtained from Lumiprobe. ATTO647N-maleimide (catalog no. AD647N-41) was obtained from ATTO-TEC. Janelia Fluor 549-NHS ester (catalog no. 6147) was obtained from Tocris. TCEP hydrochloride (catalog no. C4706-2G), dimethylformamide (DMF) (catalog no. 227056-100ML), and dithiothreitol (DTT) (catalog no. D0632-5G) were obtained from Sigma-Aldrich. Absolute ethanol (catalog no. 1009831000) was obtained from Merck. Sodium bicarbonate (catalog no. 27778293) was obtained from VWR. Tris-HCl 1 M pH 8 and 7.5, 10x and 1X PBS pH 7.3, 1X TAE pH 8, 1 M HEPES pH 7.3, EDTA 0.5 M pH 8, 1 M magnesium chloride, 1 M potassium chloride and 5 M sodium chloride were obtained from our in-house media kitchen. Pure water was obtained from a MilliQ water filtration system. SYBR Safe (catalog no. S33102) was obtained from Invitrogen. Neutravidin protein (catalog no. 31000) was obtained from Thermo Scientific. Polyethylene glycol (PEG)-8000 (catalog no. P2139-500G) and Bovine Serum Albumin-biotin (BSA-biotin) (catalog no. A8549-10MG) were obtained from Sigma-Aldrich. Round and square # 1.5 coverslips (catalog no. 6311344 and MENZBB024024AC23 respectively) were obtained from Menzel Gläser. Flat microscope slides (catalog no. 09-3000) were obtained from Bio-Optica and Sail brand single concave glass slides (catalog no. 7103) were obtained from SmartLabs. Hellmanex III (catalog no. Z805939-1EA) was obtained from Sigma-Aldrich. Double sided tape (catalog no. 665D) was obtained from Scotch. Twinsil speed 22 (catalog no. 1300-1002) was obtained from Picodent. Epoxy glue (catalog no. G14250) was obtained from Thorlabs. Tween 20 (catalog no. P9416-50ML), protocatechuate 3,4-dioxygenase from Pseudomonas sp. (PCD) (catalog no. P8279-25UN), 3,4-dihydroxybenzoic acid (PCA) (catalog no. 37580-25G-F), (+−)-6-hydroxy-2,5,7,8-tetramethylchromane-2-carboxylic acid (Trolox) (catalog no. 238813-1G), methyl viologen (MV) (catalog no. 856177-1G) and sodium hydroxide (catalog no. 30620-1KG-M) were obtained from Sigma-Aldrich. Methanol (catalog no. 20846.326) and anhydrous D-glucose (catalog no. 0188-1KG) were obtained from VWR. Glycerol (catalog no. A0970-1000) was obtained from BioChemica. Glucose oxidase (catalog no. G2133-10KU), catalase (catalog no. C100-50MG) and cysteamine hydrochloride (MEA) (catalog no. M6500-25G) were obtained from Sigma-Aldrich. 1X Dulbecco’s modified Eagle medium (DMEM) (catalog no. 11880028), 100X MEM non-essential amino acids solution (NEAA) (catalog no. 11140035), Fetal Bovine Serum (FBS) (catalog no. 10500064) and trypsin-EDTA 0.5% (catalog no. 15400054) were obtained from Gibco. L-glutamine (catalog no. 25030024) was obtained from Thermo Scientific. Penicillin-streptomycin (catalog no. P0781) was obtained from Sigma-Aldrich. The cell line is available via Cell Line Services (Nup96–SNAP, 300444). Paraformaldehyde (PFA) (catalog no. 15714) was obtained from Electron Microscopy Sciences. Triton X-100 (catalog no. T8787-100ML) was obtained from Sigma-Aldrich. Ammonium chloride (catalog no. 1011451000) was obtained from Merck. BSA (catalog no. 422351S) was obtained from VWR. Image-iT FX Signal Enhancer (catalog no. I36933) was obtained from Invitrogen. SNAP-Surface Alexa Fluor 647 (catalog no. S9136S) was obtained from NEB. 40 nm and 90 nm diameter gold nanoparticles (catalog no. G-40-100 and G-90-100) were obtained from cytodiagnostics. 100 nm Tetraspeck beads (catalog no. T7279), 40 nm dark-red beads (catalog no. F8789) and 40 nm red-orange beads (catalog no. F8794) were obtained from Invitrogen. Norland Optical Adhesive 74 (catalog no. 7404001) was obtained from APM Technica.

#### Fluorescent microspheres samples

Different fluorescent microspheres (“beads”) samples were used for calibrations, aberration correction, and testing the system. Glass coverslips were first cleaned via three rounds of water-bath sonication for 10 min in a 2% solution of Hellmanex III. Between each round and at the end, the slides were copiously rinsed with MilliQ water, and finally dried with compressed nitrogen before storage. A flow chamber was formed by placing two double-sided Scotch strips parallel to each other on the microscope slide, and then sandwiching them between the slide and the coverslip. First, 20 μl of a 1:5 dilution of 90 nm gold nanoparticles in 1X PBS was incubated for 5 min in the flow chamber. After a 20 μl wash with 1X PBS, the bead solution was incubated for 10 min. For Tetraspeck beads, this consisted of a 1:250 dilution of beads in 1X PBS, sonicated for 10 min. For the dark-red and red-orange 40 nm beads, this consisted of a 1:1000 dilution in 1X PBS, sonicated for 10 min, and then further diluted 1:1000 in 1X PBS before incubation. After a 20 μl wash with 1X PBS, the chamber was filled with more 1X PBS, and then sealed with epoxy glue.

#### Stock solutions

100X MEA: 568 mg of cysteamine hydrochloride in 5 ml of water, pH adjusted to 8 with NaOH and stored at -20°C.

100X GLOX: 2 mg of glucose oxidase in 1xPBS + a variable volume of catalase (dependent on the product batch) for a final concentration of 40 mg/ml glucose oxidase and 6.4 mg/ml catalase, stored at 4°C for a maximum of two weeks.

100X MV: 156 mg of MV in 5 mM of 50 mM Tris-HCl pH 8, stored at -20°C.

100X Trolox: 100 mg of Trolox dissolved in 430 μl of methanol, then added to 345 μl of 1 M NaOH in 3.2 ml of water, stored at -20°C.

40X PCA: 154 mg of PCA in 10 ml of water, pH adjusted to 9 with NaOH and stored at -20°C.

500X PCD: 9.3 mg of PCD in 2.66 ml of 100 mM Tris-HCl pH 8, 50 mM KCl, 1 mM EDTA, 50% glycerol, stored at -20°C.

2X TCEP: 11.5 mg of TCEP hydrochloride in 20 ml of 20 mM HEPES pH 7.3 + 100 mM NaCl, pH adjusted to 7.3 with NaOH and stored at -20°C.

#### Gold nanoparticles samples

Gold nanoparticle samples were used for PSF measurements and for the speed characterization of the axial scanner via backscattering. A flow chamber was made, and 40 nm gold nanoparticles were incubated as described above. Afterwards, the NOA 74 optical adhesive, matching the refractive index of the immersion oil, was flowed in the chamber, replacing the aqueous buffer, and cured by exposing to UV light for 15-20 minutes with a handheld UV lamp.

#### Design, folding and purification of DNA origami nanostructures

All DNA origamis are based on the p7249 scaffold derived from the M13mp18 bacteriophage. The two-dimensional rectangle origami is based on the design in^28^, while the three-dimensional 32-helix bundle (32HB) origami was designed using Cadnano2 (see Supplementary Fig.) and tested in CanDo^30^. Folding of the DNA origamis was accomplished in a 50 μl one-pot reaction consisting of 10 nM scaffold strand (20 nM for the 32HB), 100 nM unmodified staple strands, excluding the ones substituted by a modified staple strand (200 nM for the 32HB), 500 nM biotynilated staples (1 μM for the 32 HB origami) (for a list of the unmodified and biotynilated staple strands, see Supplementary Table S1 and Supplementary Table S2) and 500 nM modified staple strands (1 μM for the 32HB) (for a list of the modified staples strands, see Supplementary Table S3) in 1X folding buffer (5 mM Tris-HCl pH 8, 1 mM EDTA, 12.5 mM MgCl_2_ or 20 mM MgCl_2_ for the 32HB). The reaction mix was then subjected to a thermal annealing ramp in a thermocycler. For the rectangle origami, this consisted in 5 min at 80°C, then cooling immediately to 60°C followed by cooling from 60°C to 4°C in steps of 1°C every 3 min 21 s. The sample was then held at 4°C until purification. For the 32HB, the ramp consisted in 15’ at 65°C, then cooling immediately to 60°C followed by cooling from 60°C to 44°C in steps of 1°C every 60 min, followed by cooling from 44°C to 25°C in steps of 1°C every 10 min. The sample was then cooled immediately to 4°C and held at that temperature until purification.

In order to remove most of the excess staple strands, the rectangle origami was initially purified via one round of PEG precipitation by adding the same volume of PEG buffer (1X TAE pH 8.0 + 15% PEG-8000, 500 mM NaCl, 12.5 mM MgCl_2_ or 20 mM MgCl_2_ for the 32HB), centrifuging at 14,000 *g* at 4 °C for 30 min, removing the supernatant and resuspending in folding buffer. In the case of the 32HB, in order to select properly-folded origami, this purification was instead carried out via agarose gel electrophoresis (1.5% agarose, 0.5× TAE, 20 mM MgCl_2_, 1× SYBR Safe, run in 0.5× TAE, 20 mM MgCl_2_ as running buffer) at 3 V cm^−1^ for 3 h at 4°C. The fastest running band, corresponding to properly-folded origami, was excised on a UV transilluminator, crushed, filled into a Freeze ‘N Squeeze column, incubated for 5 min at -20°C and centrifuged for 3 min at 13000 *g* at room temperature.

After this initial purification, the origami solution was incubated with 1 μM of the appropriate fluorescent oligos (for a list of the fluorescent oligos, see Supplementary Table S4) for 2 h at room temperature, in order for it to bind to its target extended staple on the origami.

Finally, both rectangle and 32HB origamis were subjected to 3 rounds of PEG precipitation as described above to fully remove excess folding staples and fluorescent strands, and stored in low-binding Eppendorf tubes at -20°C.

#### Fluorescent labelling of oligonucleotides and purification

Oligonucleotides for external labelling of DNA origami were labelled in-house using either the maleimide + thiol or the NHS ester + amine chemistry. For the maleimide + thiol chemistry, 10 nmol of thiol-modified oligonucleotide in 50 μl of 20 mM HEPES pH 7.3 + 100 mM NaCl were added to 50 μl of 2x TCEP and incubated for 1 h at room temperature in the dark to reduce the disulfide bonds of the oligonucleotides. Then, 10 μl of 10 mM dye-maleimide in DMF were added to the reaction, the volume was brought to 200 μl with 20 mM HEPES pH 7.3 + 100 mM NaCl, and the reaction incubated for 2 h at room temperature in the dark. The reaction was then quenched by adding DTT to a final concentration of 50 mM. For the NHS ester + amine chemistry, 10 nmol of amine-modified oligos were subjected to one round of ethanol precipitation to remove any residual amine from the synthesis process by adding 0.1 volumes of 3 M NaCl and 2.5 volumes of cold absolute ethanol, incubating at - 20°C for 30 min, centrifuging at 21000 *g* at 4°C for 30 min, and resuspending the pellet in 190 μl of 100 mM sodium bicarbonate pH 8.3. Then, 10 μl of 10 mM dye-NHS ester in DMF were added to the reaction, and the reaction was incubated for 2 h at room temperature in the dark. The reaction was quenched by adding Tris-HCl pH 8 to a final concentration of 50 mM. For both chemistries, the reaction pot after quenching was subjected to one preparative round of ethanol precipitation as just described before purifying via reverse-phase high-performance liquid chromatography on an UltiMate 3000 HPLC system equipped with a Phenomenex Clarity 5 μm oligo-RP column. The purified oligos were resuspended to 50 μM in 20 mM HEPES pH 7.3 + 100 mM NaCl and stored in low-binding Eppendorf tubes at -20°C.

#### Origami sample preparation

Origami samples were prepared as previously described^19^. In short, a flow chamber was made as described above, and similarly incubated with 90 nm gold nanoparticles. The chamber was then flushed with 20 μl of 1X PBS, then with 20 μl of buffer A (10 mM Tris-HCl pH 7.5, 100 mM NaCl and 0.05% Tween 20), and then incubated with 20 μl of 1 mg ml-1 BSA-biotin in buffer A for 5 min. The chamber was then flushed with 20 μl of buffer A, and then incubated with 0.5 mg ml-1 of neutravidin in buffer A for 5 min. The chamber was then flushed with 20 μl of buffer A, then with 20 μl of buffer B (5 mM Tris-HCl pH 8, 1 mM EDTA, 12.5 mM MgCl2 and 0.05% Tween), and then incubated with 20 μl of 30 pM DNA origami in buffer B for 8 min. The chamber was then flushed with 20 μl of buffer B, and finally filled with 20 μl of the appropriate imaging buffer. For tracking experiments, this consisted of a ROXS buffer containing 1X PCA (1 mM), 1X PCD (10 nM), 1X MV (1 mM), and 1X Trolox (1 mM) in buffer B. For STORM experiments, this consisted of a blinking buffer containing 1X GLOX (0.4 mg ml-1 glucose oxidase, 64 μg ml-1 catalase), and 1-10 mM MEA in buffer B + 10% w/v glucose. The sample was then immediately sealed with Twinsil. The buffer was replaced approximately every 2 h in the case of the blinking buffer to avoid unwanted blinking behavior due to pH drift caused by the oxygen scavenging system.

#### U2OS NUP96-SNAP cell sample preparation

Labelling of the U2OS NUP96-SNAP cell samples was performed similarly to previously described^8^. Cells were cultured in DMEM without phenol red supplemented with 1X MEM NEAA, 10% v/v FBS, 2 mM L-glutamine and 1:100 penicillin-streptomycin at 37°C, 5% CO_2_ and 100% humidity and passaged every other day. The day before imaging, cells were seeded on round coverslips, cleaned as described above, so as to reach around 70% confluency the next day. The next day, cells were prefixed with 2.4% v/v PFA in 1X PBS for 30 s, permeabilized with 0.4% v/v Triton X-100 in 1X PBS for 3 min and fixed with 2.4% v/v PFA in 1X PBS for 30 min. Fixation was quenched by incubating with 100 mM ammonium chloride in 1X PBS for 5 min, followed by washing twice with 1X PBS for 5 min. The sample was blocked with Image-iT FX Signal Enhancer for 30 min, and then stained with 1 μM SNAP-Surface Alexa Fluor 647 in 1X PBS + 1 μM DTT + 0.5% w/v BSA for 2 h at room temperature. Excess unbound dye was removed by washing three times with 1X PBS for 5 min. Cells were then incubated for 10 min with 90 nm gold nanoparticles diluted 1:1 in 1X PBS, then washed three times with 1X PBS for 5 min to remove unbound nanoparticles. The sample was then mounted on a concave microscope slide, filled with blinking buffer containing 1X GLOX (0.4 mg ml^-1^ glucose oxidase, 64 μg ml^-1^ catalase), and 10-30 mM MEA in 50 mM Tris-HCl pH 8 + 10% w/v glucose, and sealed with Twinsil. The buffer was replaced approximately every 2 h to avoid unwanted blinking behavior due to pH drift caused by the oxygen scavenging system.

### MINFLUX acquisitions

The acquisition for imaging follows the previous work ^8^. For tracking, we first centered on the emitter(s) of interest in xy by moving the tip/tilt mirror controlled by a feedback loop and in z by deflecting the bottle beam. After converging to a location, the tracking took place within a “static” excitation pattern.

For detailed acquisition settings see Supplementary Note 4.

### Live localization estimator

The live estimators utilized in all MINFLUX acquisitions reported in this work are from reference ^8^, except for the one associated with the excitation scheme comprising 7 Gaussian exposures positioned at the vertices and center of a regular hexagon, which we describe below:

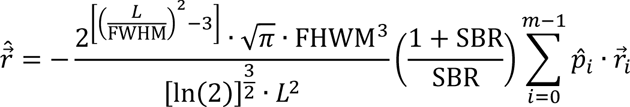

where L is the hexagon radius, FWHM is the full width at half maximum of the focused 640 nm Gaussian beam in xy (280 nm in this work), SBR is the signal-to-background ratio, 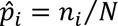 is the estimated photon arrival probability of the *i*-th exposure, and 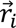 is the beam position of the *i*-th exposure. Notably, we define the SBR from the maximum and minimum photon counts within a single excitation cycle: SBR = (*n*_max_ − *n*_min_)⁄*n*_min_.

### Data analysis

All acquired MINFLUX data was processed with custom processing software implemented in Python. The final localizations were estimated by maximum likelihood estimation implemented via a gradient descent on the likelihood function. For imaging, the processing followed the previous work^8^. The background localizations were filtered as listed in Supplementary Note 5. For tracking, see the remarks below.

#### Tracking

The raw photon counts underwent binning prior to localization estimation, allowing users to specify multiple binnings and thereby tailor the trade-off between spatial and temporal resolution during processing. For reporting localization precisions, we first applied a low-pass filter to the binned counts using a rolling window, and then ran a k-means cluster algorithm on the low-pass filtered counts. Subsequently, we computed standard deviations for each cluster and averaged the two resulting numbers for x, y, and z, separately. To report the distance between binding sites in the axial oscillator, we averaged each of the two clustered traces in groups of 20 localizations and then computed standard deviations. Only the dual-color origami was drift corrected by subtracting the trend obtained when fitting a fourth order polynomial to the x, y, and z trajectories, separately.

## Extended Data

**Extended Fig. 1.**
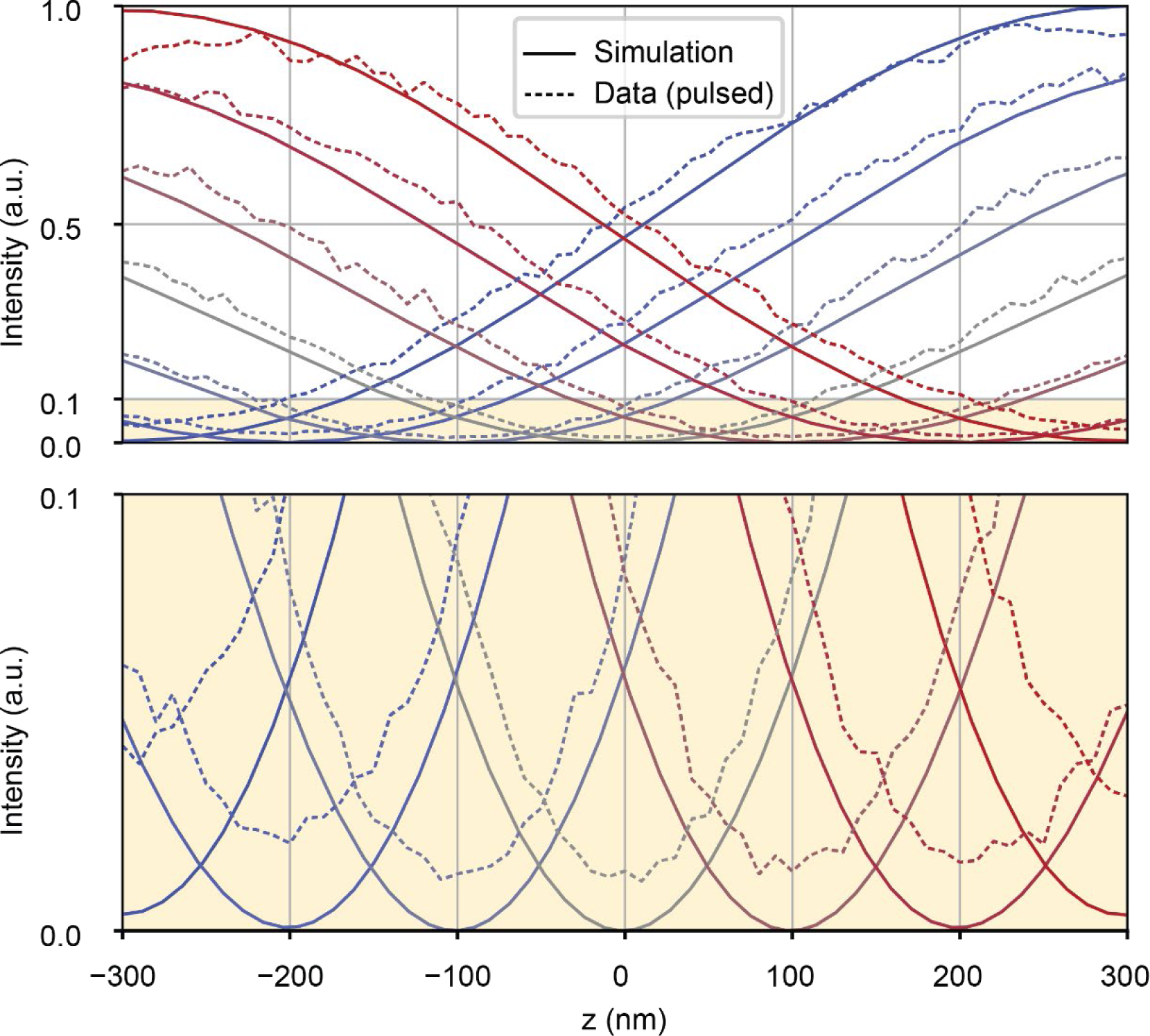
Pulsed-laser bottle beam deflection. Juxtaposition of simulation and measurement with a pulsed excitation laser of intensity profiles for different positions of the deflected bottle beam zero. Bottom: zoom-in of the yellow area. A slightly stronger zero degradation than shown in Fig. 3**c** is visible for the measured data here.

