## Supplementary Information for "A Fast Interferometric Beam Shaper for Multi-Emitter 3D MINFLUX"

### Supplementary Notes

### Supplementary Figures

### Supplementary Tables

Supplementary Table S1. Focal lengths of lenses in the system. ..... 20
Supplementary Table S. 2D rectangle origami staple strands. ..... 25
Supplementary Table S. 3D 32 HB origami staple strands. ..... 28
Supplementary Table S. Modified staples for dye attachment. ..... 31
Supplementary Table S. Fluorescent oligos. ..... 32

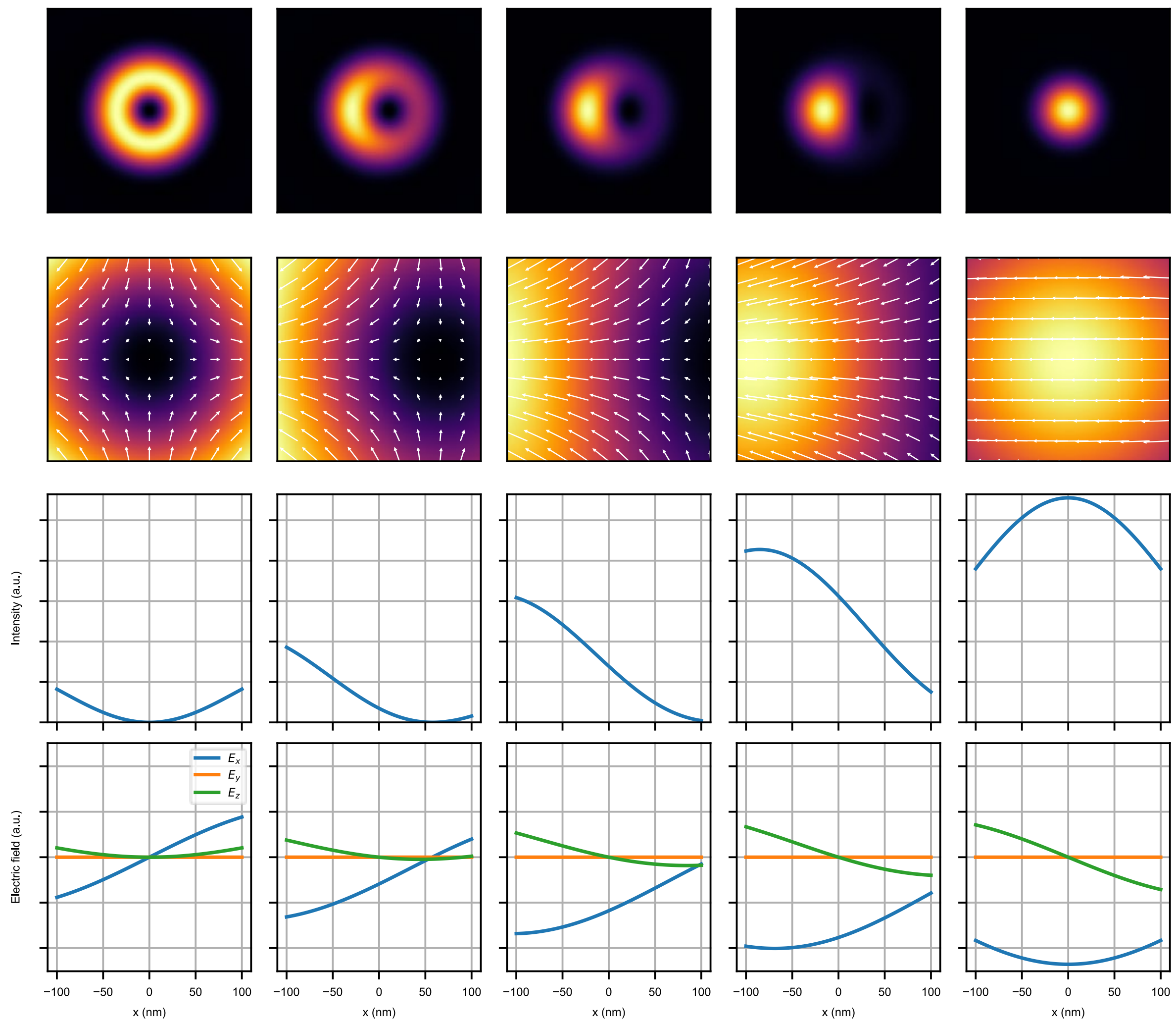

**Supplementary Fig. S1. Donut to Gaussian transition.**
PSFs for five different relative amplitudes showcasing the complete transition from a pure donut to a pure Gaussian. Top row: intensity of the xy-plane, 800x800nm. Second row: zoom-in of the xy plane, 200x200nm, where arrows indicate the electric field components. Last two rows: Intensity and electric field profiles as labelled.

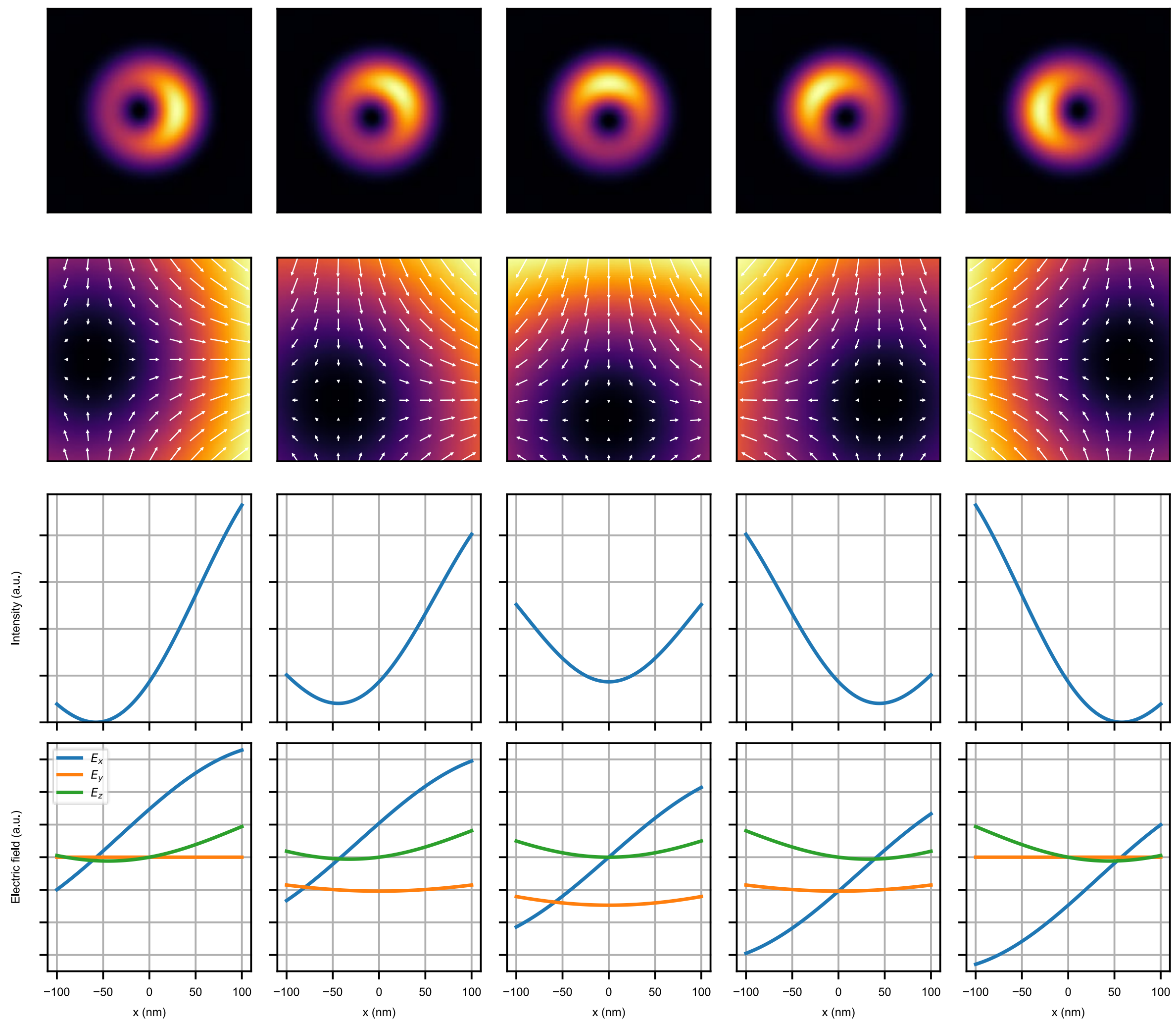

**Supplementary Fig. S2. Donut rotation.**
PSFs of a donut plus Gaussian combination for five different relative phases showcasing a 180° rotation of the zero. Top row: intensity of the xy-plane, 800x800nm. Second row: zoom-in of the xy-plane, 200x200nm, where arrows indicate the electric field components. Last two rows: Intensity and electric field profiles as labelled.

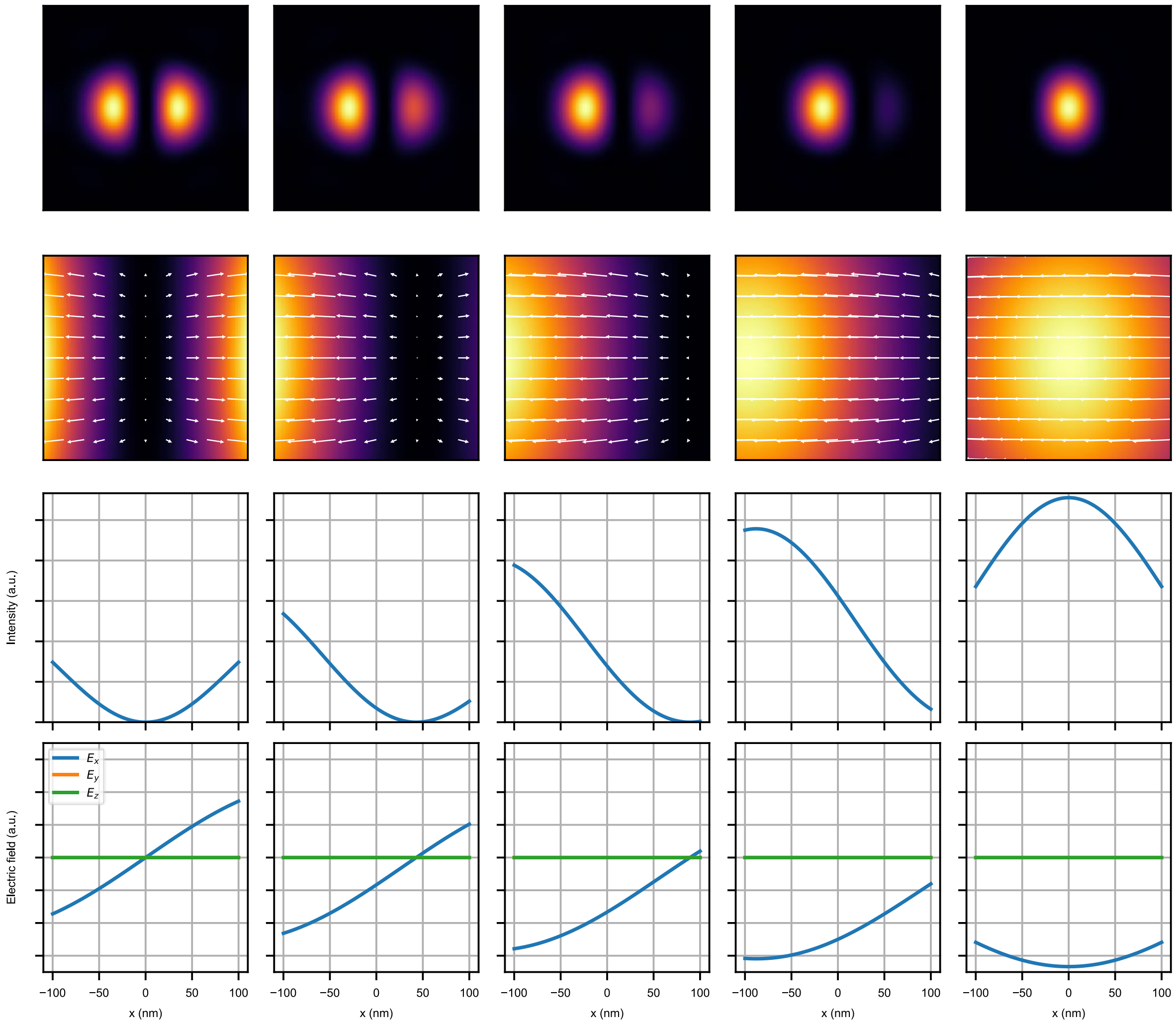

**Supplementary Fig. S3. Bi-lobed to Gaussian transition.**
PSFs for five different relative amplitudes showcasing the complete transition from a pure bi-lobed beam to a pure Gaussian. Top row: intensity of the xy-plane, 800x800nm. Second row: zoom-in of the xy plane, 200x200nm, where arrows indicate the electric field components. Last two rows: Intensity and electric field profiles as labelled.

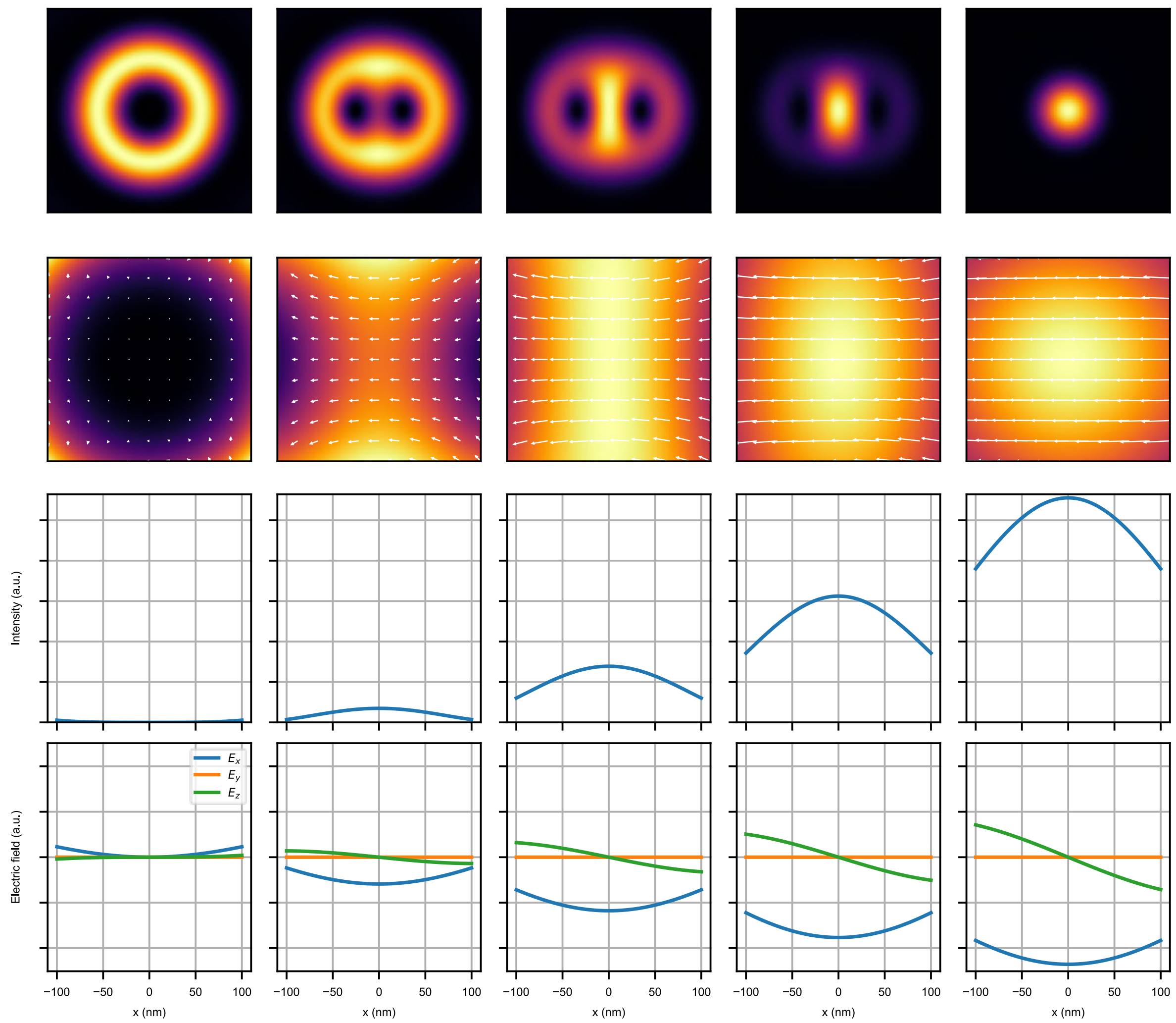

**Supplementary Fig. S4**. **Charge 2 donut to Gaussian transition.**
PSFs for five different relative amplitudes showcasing the complete transition from a pure charge 2 donut to a pure Gaussian. Top row: intensity of the xy-plane, 800x800nm. Second row: zoom-in of the xy-plane, 200x200nm, where arrows indicate the electric field components. Last two rows: Intensity and electric field profiles as labelled.

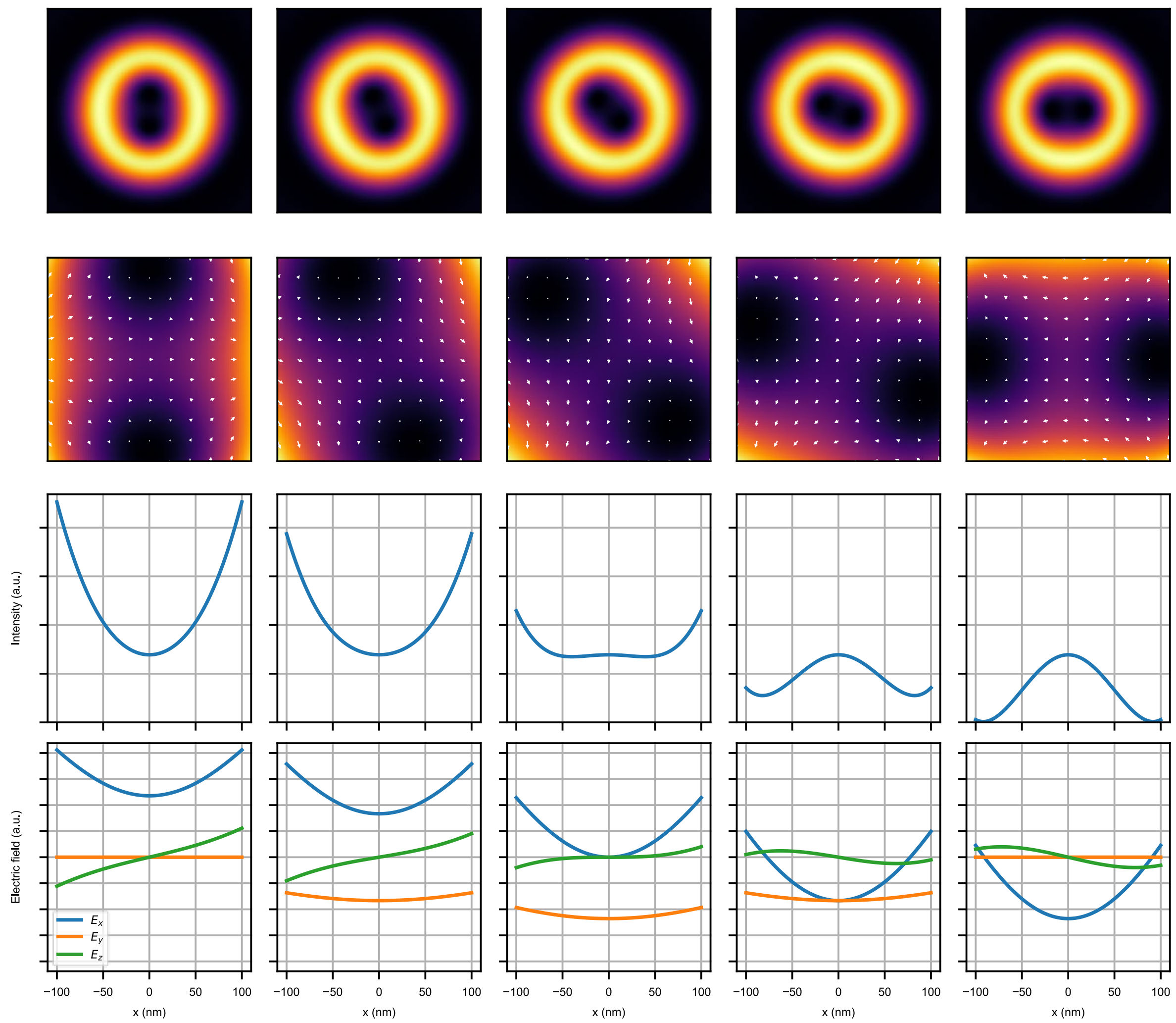

**Supplementary Fig. S5. Charge 2 donut rotation.**
PSFs of a charge 2 donut plus Gaussian combination for five different relative phases showcasing a 90° rotation of two zeros. Top row: intensity of the xy-plane, 800x800nm. Second row: zoom-in of the xy-plane, 200x200nm, where arrows indicate the electric field components. Last two rows: Intensity and electric field profiles as labelled.

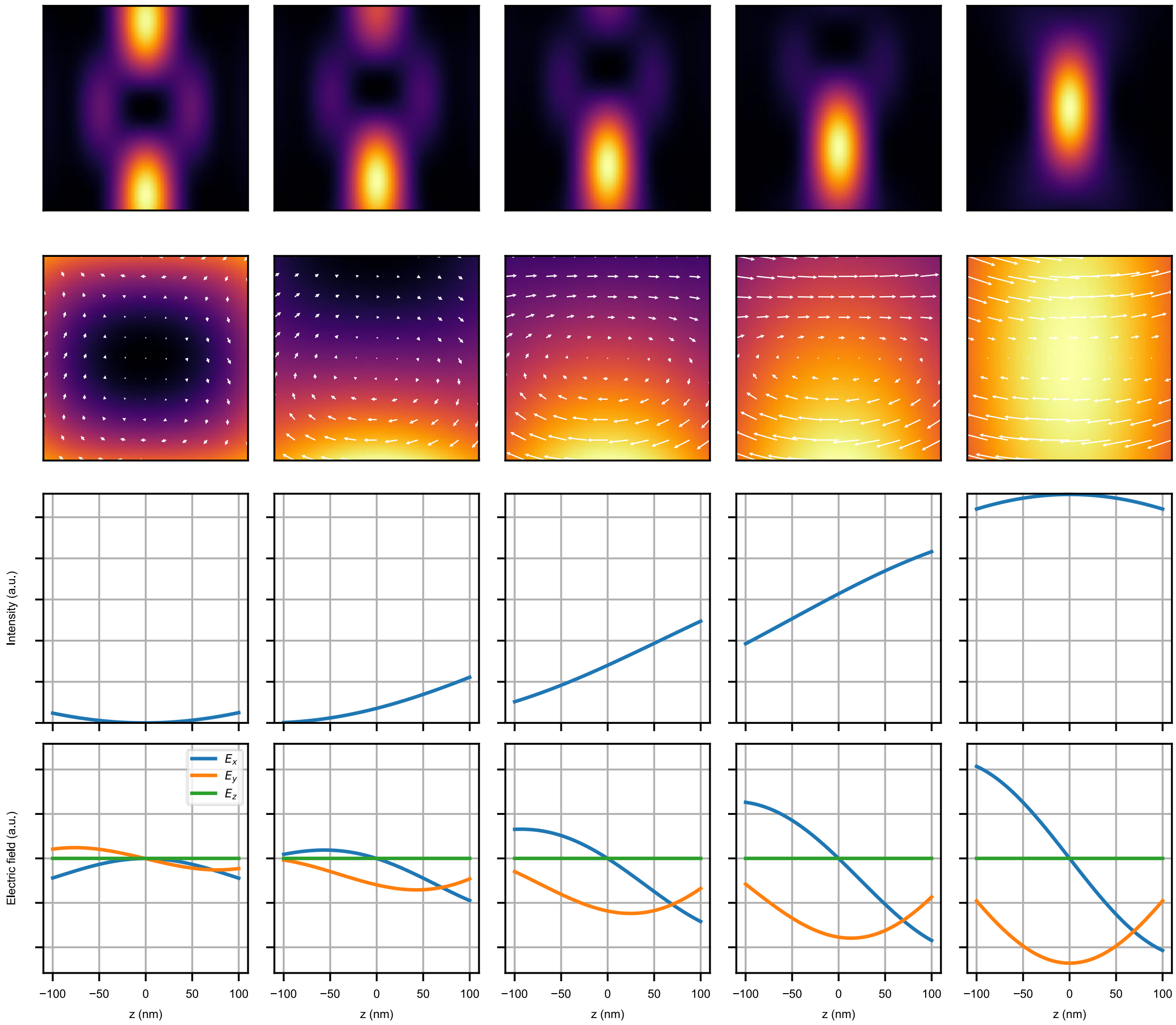

**Supplementary Fig. S6**. **Bottle beam to Gaussian transition.**
PSFs for five different relative amplitudes showcasing the complete transition from a pure donut to a pure Gaussian. Top row: intensity of the xy-plane, 800x800nm. Second row: zoom-in of the xy plane, 200x200nm, where arrows indicate the electric field components. Last two rows: Intensity and electric field profiles as labelled.

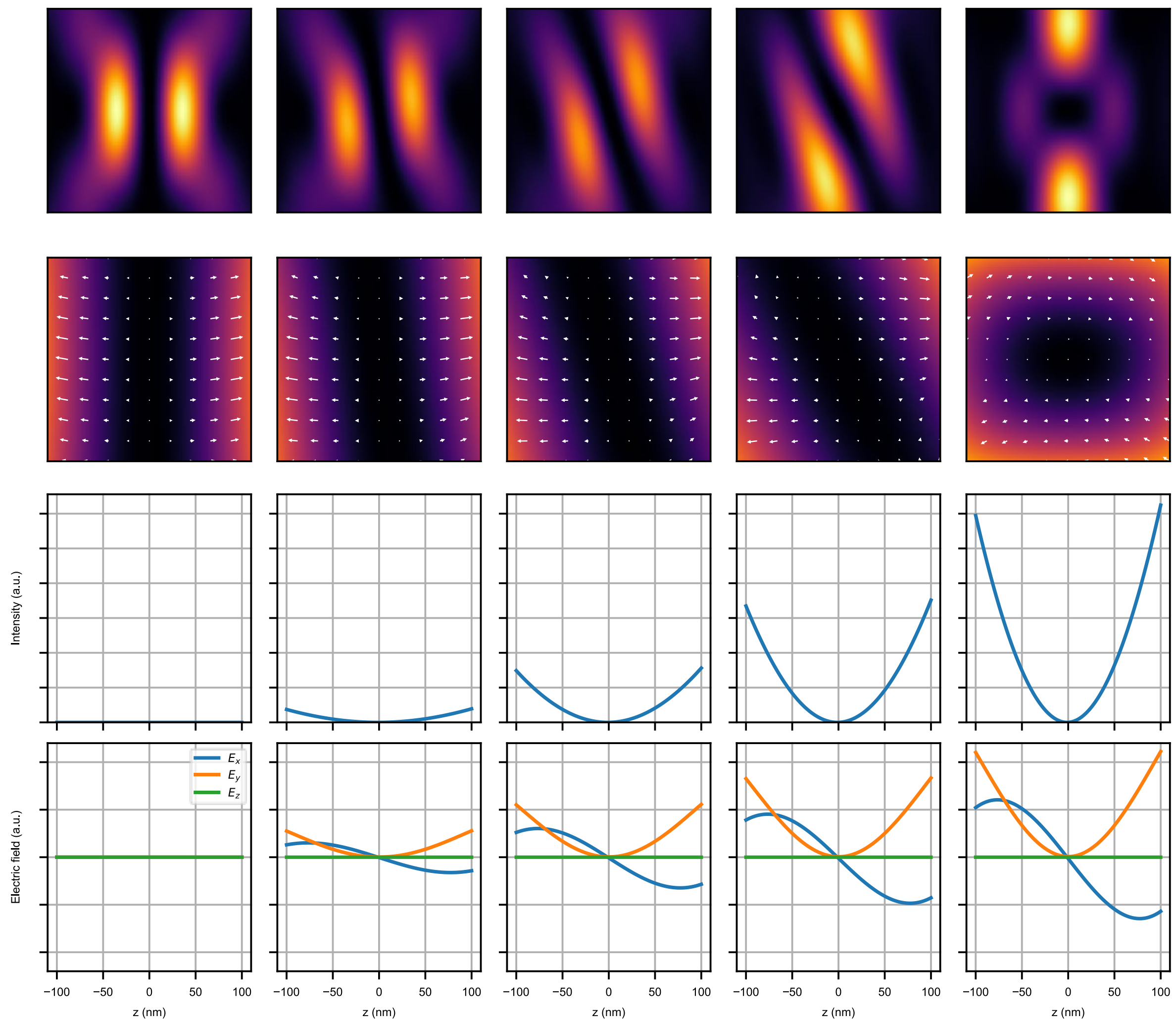

**Supplementary Fig. S7**. **Donut to bottle beam transition.**
PSFs for five different relative amplitudes showcasing the complete transition from a pure donut to a pure bottle beam. Top row: intensity of the xz-plane, 1200x1200nm. Second row: zoom-in of the xz plane, 200x200nm, where arrows indicate the electric field components. Last two rows: Intensity and electric field profiles as labelled.

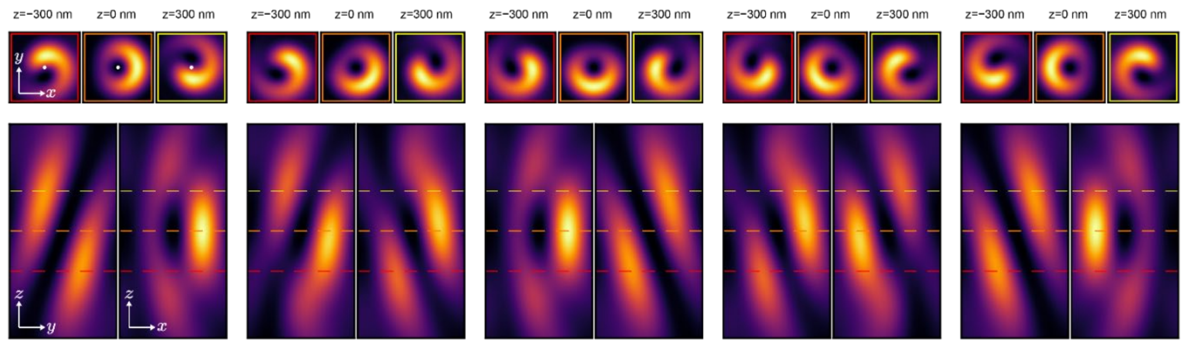

**Supplementary Fig. S8. Donut tilt-axis rotation.**
PSFs of a donut-plus-bottle-beam combination for five different relative phases showcasing a 180° rotation of the tilted zero axis. Top row: intensity of the xy-plane, 800x800nm, at different values of z. The colored frames indicate the z coordinate (red: -300 nm, orange: 0 nm, yellow: +300 nm). The white dots in the first three plots mark the center of the PSF (i.e., the position of the zero for z = 0 nm) to highlight that the tilt axis is oriented along y. While the tilt does not move the zero at z = 0 nm, the PSF showcases an asymmetric drop in intensity lateral to the tilt plane (i.e., in along x) resembling the PSF of a donut with a deflected zero. Second row: yz- and xz-planes, 800x1600nm. The colored dashed lines indicate the z coordinates of the xy planes as described above.

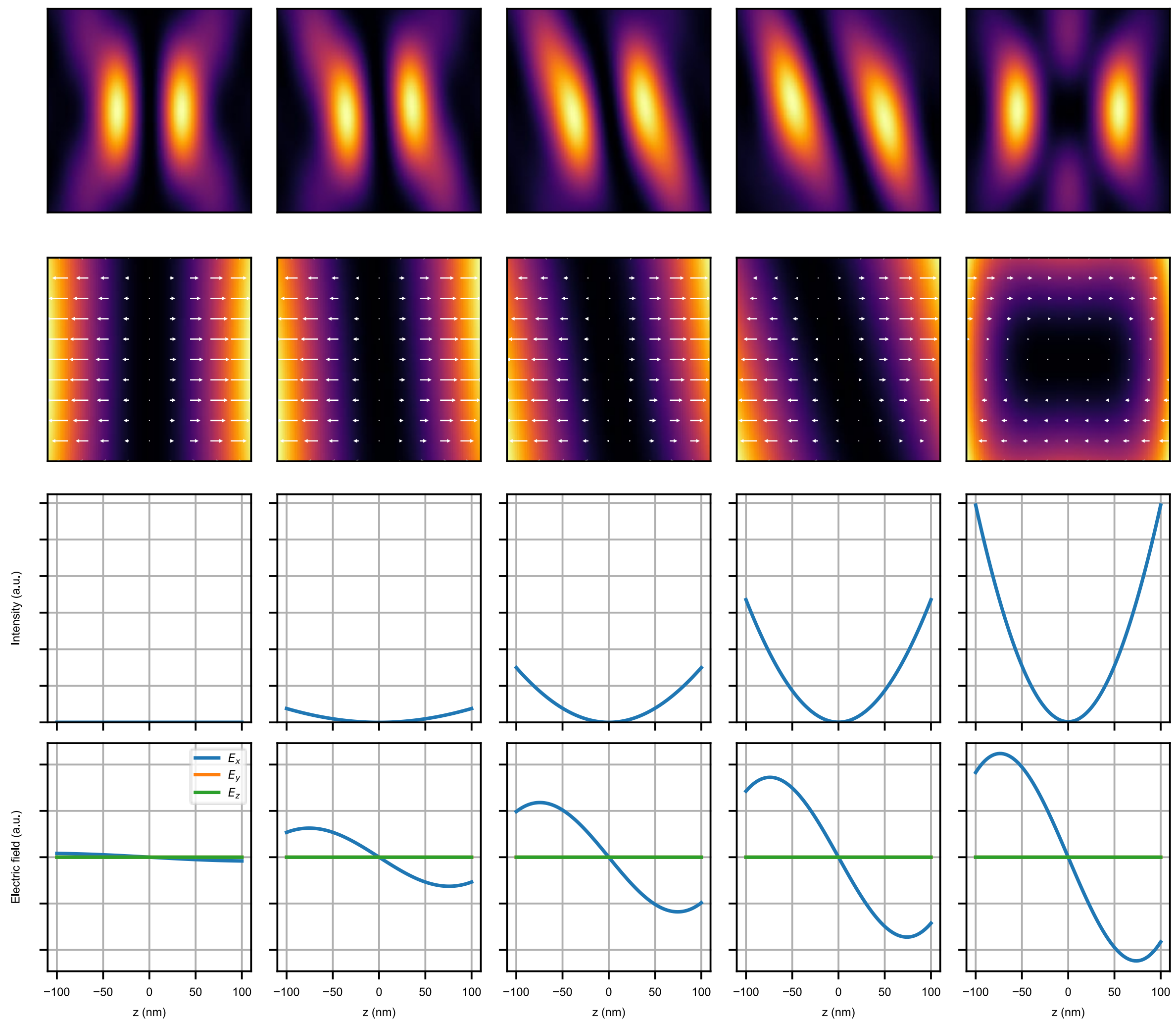

**Supplementary Fig. S9**. **Bi-lobed to bottle-slice beam transition.**
PSFs for five different relative amplitudes showcasing the complete transition from a pure bi-lobed to a pure bottle-slice beam. Top row: intensity of the xz-plane, 1200x1200nm. Second row: zoom-in of the xz plane, 200x200nm, where arrows indicate the electric field components. Last two rows: Intensity and electric field profiles as labelled.

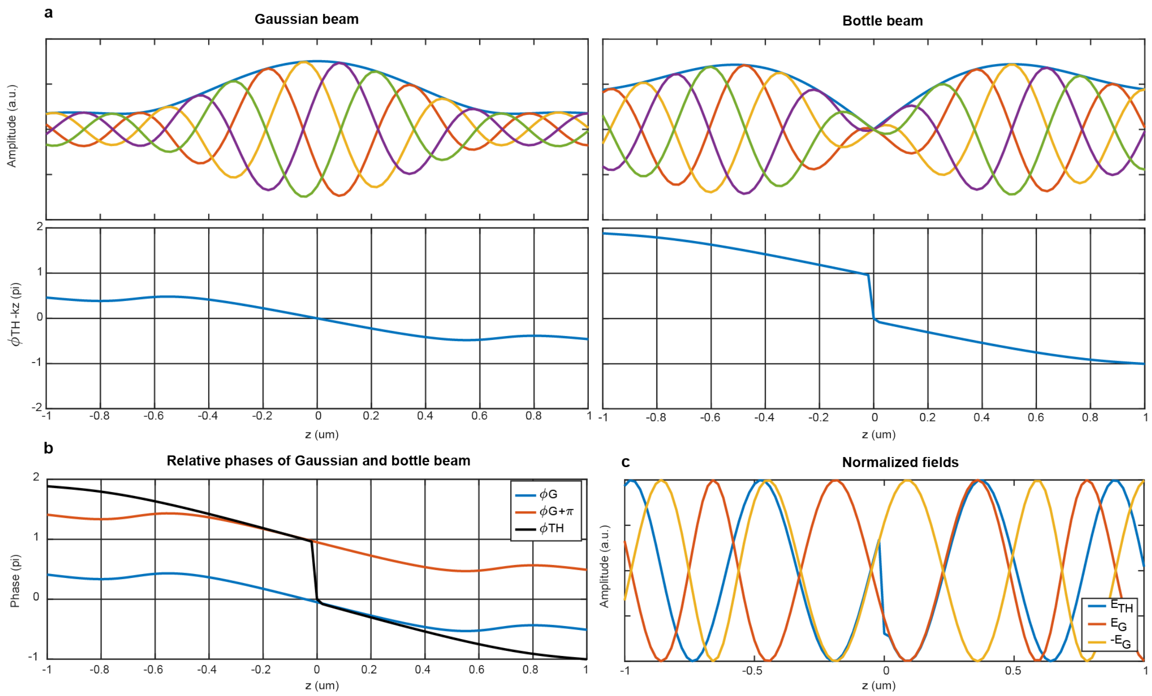

**Supplementary Fig. S10. Gouy phase of Gaussian and bottle beam.**

**a**, Top row: Real part of the electric field along z for the Gaussian and the bottle beam for several time points (colored lines) and the absolute value (blue envelope). Bottom row: Gouy phase of the Gaussian and bottle beam. While the Gaussian beam smoothly acquires a π phase shift, the bottle beam acquires 2π plus an additional π (phase jump) at the zero. b, Gouy phase of the Gaussian with and without a π offset (colored lines) and the bottle beam (black). The mismatch grows resulting in the degradation of the bottle beam zero when deflected. c, Real parts of the normalized fields of the Gaussian (orange and yellow) and the bottle beam (blue). While they propagate together close to the focus, they lose synchronization farther off, leading to zero degradation.

### Supplementary Note 1. SLM double pass Jones model

In this section we develop a model that accounts for imperfections of optical elements involved in the SLM double pass configuration and distinguishes the polarization state of distinct diffraction orders involved.

The first telescope after the Spatial Light Modulator (SLM) double pass contains an adjustable iris acting like a pinhole of a spatial filter, however, it is set up to only allow the first order to pass. It is not only supposed to block the zero-th order but also other orders, which contain polarization crosstalk.

For instance, a perfectly vertical input state is patterned on the first SLM pass with a diffraction grating. Hence, the order selector would let it pass. However, if the polarization is not perfectly flipped to the horizontal state after going through the quarter-wave plate (QWP) twice, the residual vertical state is patterned on the second SLM pass. Since the second pass is showing a grating again, the residual state acquires an extra diffraction angle and will not pass the order selector. The same applies to a perfectly horizontal input state and hence any mixture. This circumstance highlights the importance of having a clean input state in cases where only a single pass is to be addressed; the polarization crosstalk of all other imperfections is automatically filtered out. Different configurations of off/on states of the gratings on the first and second pass allow to isolate these crosstalk components which is useful for characterization.

With these observations, we model the double pass with an extended Jones formalism:

$$\vec{J}^{\text{out}} = \text{PBS}\ \text{HWP}(\theta_{\text{HWP}})\ \text{SLM}_2\ \widetilde{\text{QWP}}(\text{-}\theta_{\text{QWP}}, \Delta\psi)\ \tilde{\text{L}}\ \tilde{\text{M}}\ \tilde{\text{L}}\ \widetilde{\text{QWP}}(\theta_{\text{QWP}}, \Delta\psi)\ \text{SLM}_1 \vec{J}^{\text{in}} \qquad \textbf{(1)}$$

where:

| | |
|---|---|
| $\vec{J}^{\text{in}} = \begin{pmatrix} \sin(\alpha)\, E_2 \\ \cos(\alpha)\, E_1 e^{i\Delta\varphi_{\text{sp}}} \end{pmatrix}$ | Input Jones vector. |
| $E_1$ | Input electric field amplitude, s-polarization. |
| $E_2$ | Input electric field amplitude, p-polarization. |
| $\alpha$ | Rotation angle of linear input polarization ($\alpha = 0°$ is vertical). |
| $\Delta\varphi_{\text{sp}}$ | Relative phase between s- and p-polarization. |
| $\text{SLM}_1 = \begin{pmatrix} 1 & 0 \\ 0 & \varepsilon \\ 0 & 0 \\ 0 & (1-\varepsilon)e^{i\varphi_1(x,y)} \end{pmatrix}$ | Extended Jones matrix for the first SLM pass. |
| $\varepsilon$ | Fraction of vertically polarized light that is not diffracted by the SLM. In the following we assume $\varepsilon = 0$. |
| $\text{SLM}_2 = \begin{pmatrix} 0 & 0 & 1 & 0 \\ 0 & e^{i\varphi_2(x,y)} & 0 & 0 \end{pmatrix}$ | Extended Jones matrix for the second SLM pass. |
| $\varphi_1(x,y)$ | Phase mask of first pass. |
| $\varphi_2(x,y)$ | Phase mask of second pass. |
| $\text{QWP}(\theta_{\text{QWP}}, \Delta\psi) = R(\theta_{\text{QWP}}) \begin{pmatrix} 1 & 0 \\ 0 & -i\, e^{i\Delta\psi} \end{pmatrix} R^{-1}(\theta_{\text{QWP}})$ | Jones matrix of an imperfect quarter-wave plate. |
| $\theta_{\text{QWP}}$ | Rotation angle of the quarter-wave plate |

| | |
|---|---|
| $\Delta\psi_{\mathrm{QWP}}$ | Deviation from the ideal retardation of the quarter-wave plate. |
| $R(\theta) = \begin{pmatrix} \cos(\theta) & -\sin(\theta) \\ \sin(\theta) & \cos(\theta) \end{pmatrix}$ | Rotation matrix. |
| $L = \begin{pmatrix} 1 & 0 \\ 0 & 1 \end{pmatrix}$ | Jones matrix of the lens. In our model we assume it has no effect. |
| $M = \begin{pmatrix} 1 & 0 \\ 0 & -1 \end{pmatrix}$ | Jones matrix for the metallic mirror. In our model we neglect the angle of incidence. |
| $\mathrm{HWP}(\theta_{\mathrm{HWP}}) = R(\theta_{\mathrm{HWP}}) \begin{pmatrix} 1 & 0 \\ 0 & -1 \end{pmatrix} R^{-1}(\theta_{\mathrm{HWP}})$ | Jones matrix for an ideal half-wave plate. |
| $\theta_{\mathrm{HWP}}$ | Rotation angle of the half-wave plate. |
| $\mathrm{PBS} = \begin{pmatrix} 0 & 0 \\ 0 & 1 \end{pmatrix}$ | Jones matrix for a perfect polarizer. |
| $\tilde{X} = \mathbb{1} \otimes X = \begin{pmatrix} X & \mathbb{0} \\ \mathbb{0} & X \end{pmatrix}$ | The $\tilde{}$ extends a Jones matrix via the Kronecker product $\otimes$ as described below. |
| $\mathbb{1} = \begin{pmatrix} 1 & 0 \\ 0 & 1 \end{pmatrix}$ | 2x2 unit matrix. |
| $\mathbb{0} = \begin{pmatrix} 0 & 0 \\ 0 & 0 \end{pmatrix}$ | 2x2 zero matrix. |

In essence, the model follows the conventional Jones formalism where the 2x2 Jones matrices of all optical elements are multiplied together to yield the final 2x2 matrix that acts on the input Jones vector (multiplied from the right). However, because the mirror reverts the propagation direction, the angle of the of QWP must be flipped for its second pass. Furthermore, as the SLM-passes use blazed gratings acting on the vertical polarization, the vertically polarized component of an incoming beam is

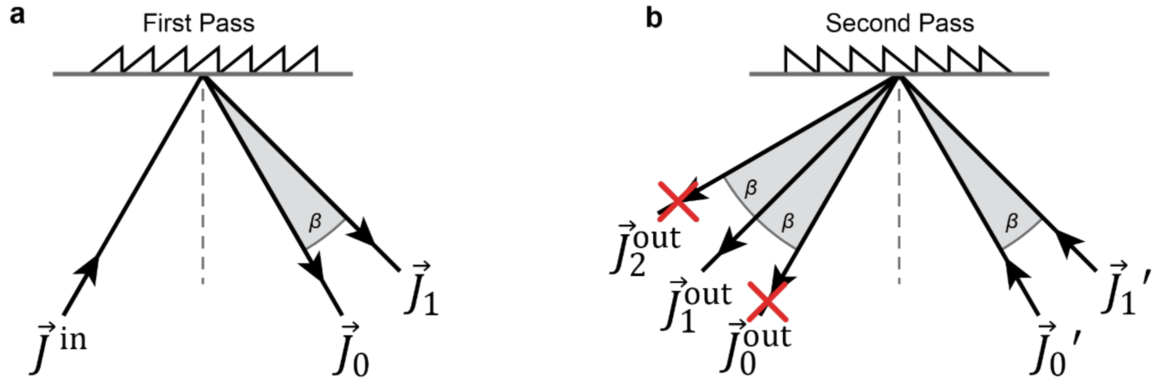

**Supplementary Fig. S11. SLM Double pass Jones vectors.**
**a**, The grating on the first SLM pass diffracts the vertically polarized component of the input beam, described by the Jones vector $\vec{J}^{\,\mathrm{in}}$, into a first order, $\vec{J}_1$, at an angle $\beta$ from the direct reflection (zero-th order), $\vec{J}_0$. **b**, After traversing the SLM double pass elements (QWP, lens, mirror) the two orders from the first pass are incident on the second pass with modified Jones vectors $\vec{J}_0{}'$ and $\vec{J}_1{}'$. Again, the vertically polarized components will be diffracted acquiring an angle of $\beta$. The horizontally polarized component of $\vec{J}_1{}'$, that already acquired the diffraction angle on the first pass, will be reflected into the beam $\vec{J}_1^{\,\mathrm{out}}$. Any “spurious” vertical component, i.e., polarization crosstalk caused by the optical elements between the passes, will be diffracted into the second order $\vec{J}_2^{\,\mathrm{out}}$. The same holds for $\vec{J}_0{}'$ but the crosstalk component will go to $\vec{J}_0^{\,\mathrm{out}}$ instead. Since the order selector (OS.PH) will block $\vec{J}_0^{\,\mathrm{out}}$ and $\vec{J}_2^{\,\mathrm{out}}$ as indicated by the red crosses, we end up with $\vec{J}_1^{\,\mathrm{out}}$ which is free of any crosstalk component.

diffracted into the first order while the horizontally polarized component is directly reflected (zero-th order). Hence, after the first pass, there are two beams each with a corresponding Jones vector $\vec{J}_0$ and $\vec{J}_1$ (Supplementary Fig. S11a), which we "concatenate" into one extended 4x1 Jones vector:

$$\begin{pmatrix} \vec{J}_0 \\ \vec{J}_1 \end{pmatrix} = \mathrm{SLM}_1\, \vec{J}^{\,\mathrm{in}}$$

Therefore, $\mathrm{SLM}_1$ is a 4x2 matrix generating the two Jones vectors. We assume that both diffraction orders are equally affected as they propagate through the optical elements between the SLM passes (QWP, lens and mirror). The matrices of these elements require an extension from 2x2 to 4x4 block matrices $X \rightarrow \begin{pmatrix} X & \mathbb{0} \\ \mathbb{0} & X \end{pmatrix}$. After the second pass, both diffraction orders diffract again, however, two of the resulting orders overlap, yielding 3 beams: $\vec{J}_0^{\mathrm{out}}$, $\vec{J}_1^{\mathrm{out}}$ and $\vec{J}_2^{\mathrm{out}}$, see Supplementary Fig. S11b.

As only $\vec{J}_1^{\mathrm{out}}$ can pass through the order selector, and there is no interaction of the orders after the SLM, our model only keeps track of $\vec{J}_1^{\mathrm{out}}$ after the second pass of the SLM. This is why we restrict $\mathrm{SLM}_2$ to a 2x4 matrix coming from the full expression:

$$\begin{pmatrix} \cancel{\vec{J}_0^{\mathrm{out}}} \\ \vec{J}_1^{\mathrm{out}} \\ \cancel{\vec{J}_2^{\mathrm{out}}} \end{pmatrix} = \mathrm{SLM}_2 \begin{pmatrix} \vec{J}_0{}' \\ \vec{J}_1{}' \end{pmatrix} = \begin{pmatrix} 1 & 0 & 0 & 0 \\ 0 & \varepsilon & 0 & 0 \\ 0 & 0 & 1 & 0 \\ 0 & (1-\varepsilon)e^{i\varphi_2(x,y)} & 0 & 0 \\ 0 & 0 & 0 & 0 \\ 0 & 0 & 0 & 1 \end{pmatrix} \begin{pmatrix} \vec{J}_0{}' \\ \vec{J}_1{}' \end{pmatrix}$$

Evaluating Eq. (1) (with $\varepsilon = 0$) yields the following output in the vertical polarization state after the PBS:

$$\vec{J}_1^{\,\mathrm{out},(1)} = \left(\sin(2\theta_{\mathrm{QWP}})\cos\left(\Delta\psi_{\mathrm{QWP}}\right)\right)\left(E_1 e^{i\,\varphi_1(x,y)}\, e^{i\Delta\varphi_{\mathrm{sp}}}\cos(\alpha)\cos(2\theta_{\mathrm{HWP}}) - E_2 e^{i\varphi_2(x,y)}\sin(\alpha)\sin(2\theta_{\mathrm{HWP}})\right)$$

The model shows that a non-ideal rotation and/or retardation of the quarter-wave plate only causes an overall loss in intensity. Furthermore, if the input is elliptical, i.e., $\Delta\varphi_{\mathrm{sp}} \neq 0$ (caused, e.g., by the reflections on mirrors following the polarization rotator) the phase difference can be compensated by the phase mask $\varphi_1$ (or $\varphi_2$) through $\varphi_1(x,y) = \tilde{\varphi}_1(x,y) - \Delta\varphi_{\mathrm{sp}}$; assuming that $\Delta\varphi_{\mathrm{sp}}$ is not a function of $\alpha$. The same argument holds for any static phase difference induced by the lens or the mirror (not modelled here).

We also note that the QWP at the polarization rotator is optional, as the EOM alone will continuously modulate the s- and p-polarization amplitudes. There is, however, a constant difference in phase of π/2 instead 0. This is equivalent to setting $\Delta\varphi_{\mathrm{sp}} = \pi/2$ if no other imperfections alter the polarization of the input $E_{\mathrm{in}}$ and can also be compensated by SLM if necessary.

For $E_0 = E_1 = E_2$, $\Delta\varphi_{\mathrm{sp}} = 0$, $\theta_{\mathrm{QWP}} = 45°$ , $\theta_{\mathrm{HWP}} = 45°/2$ we get:

$$E_{\mathrm{out}}^1 = \frac{E_0}{\sqrt{2}}\left(e^{i\,\varphi_1(x,y)}\cos(\alpha) + e^{i\varphi_2(x,y)}\sin(\alpha)\right)$$

By choosing $\varphi_2(x,y) = {}^{\pi}/_{2}$ and $\varphi_1(x,y) = \begin{cases} \pi, & (x,y) \in \mathcal{D}_1 \\ 0, & (x,y) \in \mathcal{D}_2 \end{cases}$ , where $\mathcal{D}_1$and $\mathcal{D}_2$ are two non-overlapping domains of the phase mask, we arrive at

$$E_{\mathrm{out}}^1 = \frac{E_0}{\sqrt{2}} \begin{cases} -e^{-i\,\alpha}, (x,y) \in \mathcal{D}_1 \\ \;\; e^{+i\,\alpha}, (x,y) \in \mathcal{D}_2 \end{cases}$$

This means that in $\mathcal{D}_1/\mathcal{D}_2$ we have a phasor rotating clockwise/counterclockwise depending on the angle $\alpha$ (that linearly depends on the voltage applied to the polarization rotator). The amplitude stays constant for any $\alpha$. The relative phase between the two domains is $\Delta\varphi_{\mathcal{D}} = \pi - 2\alpha$. This derivation reveals another interpretation of how the zero (point, line or plane) of any beam shape that is generated by a flat phase mask with a $\pi$ phase jump is deflected. Namely, by increasing or decreasing the magnitude of the jump. For example, an undeflected bottle beam corresponds to the "top-hat" mask with a $\pi$ phase step where $\mathcal{D}_1 = \{(x, y) | \sqrt{x^2 + y^2} < r_{\text{top-hat}}\}$. Following the above scheme, this phase step is effectively made smaller or larger when the voltage of the rotation EOM is increased or decreased, see Fig.S5. The corresponding beam shapes are deformed bottle beams with deflected zeros.

Finally, a note on the rotation angle of the half-wave plate before the interference PBS: it is not required to be at 22.5°. Changing the angle will change the relative amplitude of the two polarization states in the interference affecting the deflection range and the zero degradation. Most notably, however, the deflected distance becomes increasingly nonlinear for other half-wave plate rotation angles, see Supplementary Fig. S14.

### Supplementary Note 2: Coherence

Both polarization states must not acquire a relative retardation that is larger than the coherence length of the excitation light for interference to occur at the polarizing beam splitter after the SLM. However, during propagation, these states accumulate relative phase at every mirror following the polarization rotator.

We achieved an acceptable zero degradation with the diode laser, iBEAM-SMART-640 Ver.: G1 (Nov./2015) (TOPTICA Photonics AG, Munich, Germany). According to the manufacturer, the coherence length of this particular laser is around 4 mm with a spectral bandwidth of 0.11 nm while newer iterations of the same model have a reduced coherence length of about 1.3 mm to 0.8 mm (spectral bandwidth: 0.3 nm to 0.5 nm). We also achieved acceptable performance with an LDH-IB-640-B (PicoQuant, Berlin, Germany) in continuous wave (Fig. 3**c**) as well as pulsed operation (Supplementary Figure: pulsed laser bottle beam deflection). We did not achieve acceptable deflection with an Omicron QuixX® 642-140 (Omicron, Rodgau-Dudenhofen, Germany) diode laser, which exhibits poorer coherence.

For the green excitation we utilized Cobolt Jive™ 100-561 (Cobolt AB, Solna, Sweden), with a high coherence length of meters, hence excellent deflection performance.

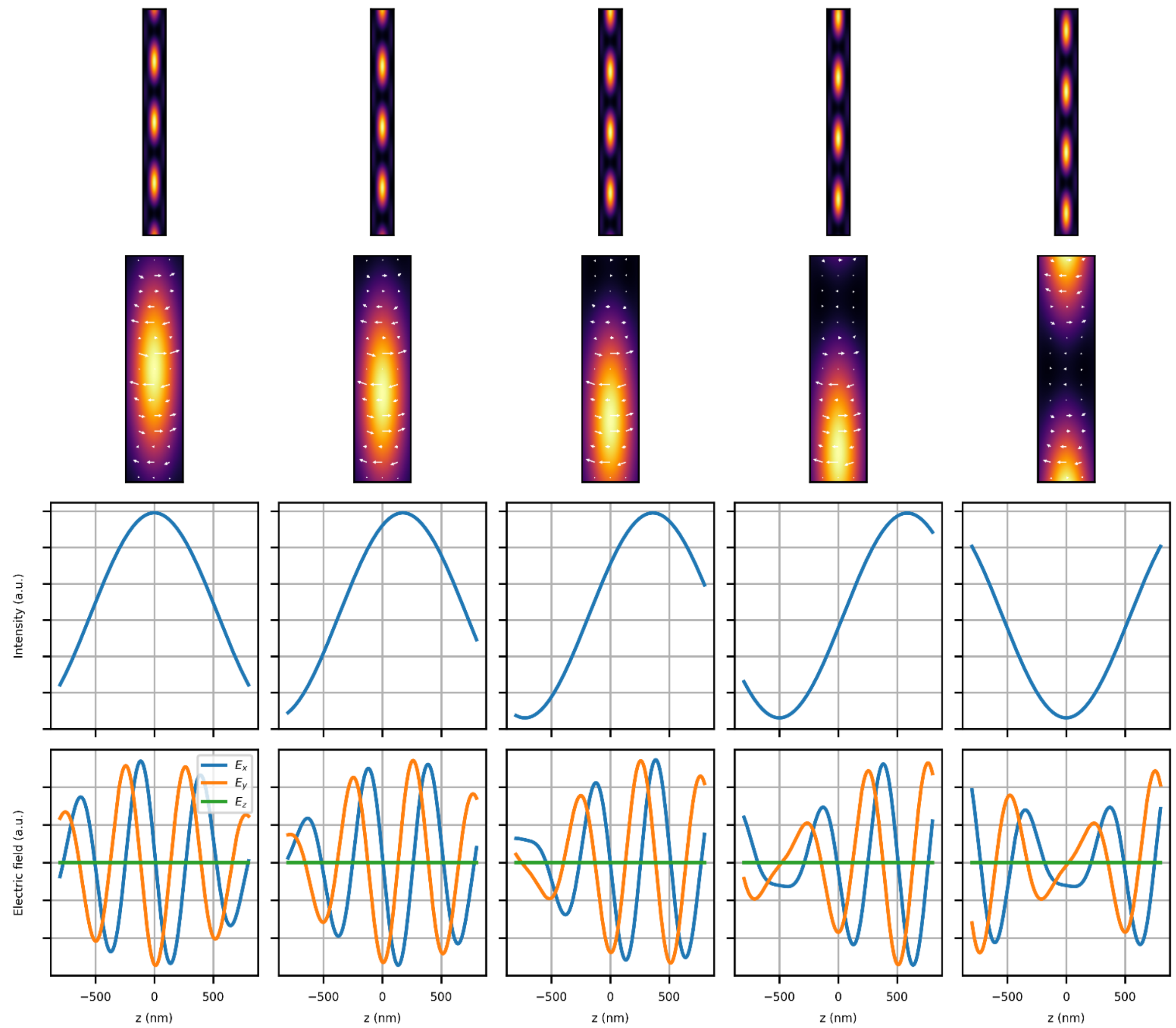

**Supplementary Fig. S12. Axial Bessel droplet scan.**
A phase mask where the blazed grating has the form of two annuli (Here: radius one 36%, radius two 71%, thickness 2% of circular SLM pupil) results in the Bessel droplet; an elongated beam with suppressed side lobes in xy[21]. Interfering the Bessel droplet beam with another Bessel droplet generated by a phase mask where the relative phase of the annuli is $\pi$, leads to a continuous scan of the droplets' positions; effectively the relative phase of the annuli is scanned, see Supplementary Fig. S15. The complete transition for five different amplitudes is shown above. Top row: intensity of the xz-plane, 0.8x8 µm. Second row: zoom-in of the xz plane, 0.4x1.6 µm, where arrows indicate the electric field components. Last two rows: Intensity and electric field profiles as labelled.

### Supplementary Note 3. Optical System

Supplementary Fig. S13 shows the detailed optical system. Mirrors lacking labels are either half-,1- or 2-inch in size and feature a visible dielectric coating, except for those in the stabilization modules, which are coated with an infrared dielectric coating.

The laser used for active stabilization in the z-axis experienced mode hopping, leading to sudden axial jumps of the fine stage. To overcome this issue, we added a diffraction grating (GR13-1210 from Thorlabs Inc., Newton, NJ, USA) mounted in the Littrow configuration. This configuration provides narrow spectral feedback to the laser diode, resulting in improved mode stability.

The addition half-wave plate A.HWP (on a rotation mount) on path A was necessary for slight adjustments of the polarization state to reduce the polarization crosstalk that was otherwise worse than that of path B.

#### List of components

**Lasers**

- E.Laser1: LDH-IB-640-B (PicoQuant, Berlin, Germany)
- E.Laser2: Cobolt Jive™ 100-561 (Cobolt AB, Solna, Sweden)
- WFE.Laser1: Omicron QuixX® 642-140 (Omicron, Rodgau-Dudenhofen, Germany)
- WFE.Laser2: Cobolt Jive™ 100-561 (Cobolt AB, Solna, Sweden)
- WFE.Laser3: iBEAM-SMART-488-S-HP (Toptica, Munich, Germany)
- AC.Laser: LBX-405 (Oxxius, Lannion, France)
- zS.Laser: MDL-XS-940 (CNI, Changchun, China)
- SLED: EXS210153-01 (Exalos AG, Schlieren, Switzerland)

**Beam modulation**

- E.AOTF: AOTFnC 400.650-CPCh-TN (AA Opto Electronic, Orsay, France)
- E.EOM: EM200K (Leysop Ltd., Essex, England) + A-304 high-voltage amplifier (A.A.Lab Systems Ltd., Ramat Gan, Israel)
- A.EOM, B.EOM: LM 0202 ADP (Excelitas Technologies, Waltham, MA, USA) + A-304 high-voltage amplifier (A.A.Lab Systems Ltd., Ramat Gan, Israel)
- DP.SLM: 1920 x 1152 Spatial Light Modulator (Meadowlark Optics, Inc., Frederick, CO, USA)

**Scanning**

- EOD: 2 mm aperture Electro-Optical Deflector (Leysop Ltd., Essex, England) + WMA-300 high-voltage amplifier + WMA-IB-HS phase inverter (Falco Systems BV, Katwijk aan Zee, The Netherlands)
- STG.TM: PSH 10/2 + NV 200/D controller (piezosystem jena GmbH, Jena, Germany)
- Coarse Stage Z: L-505.013212F (Physik Instrumente GmbH & Co. KG, Karlsruhe, Germany)
- Stage (Physik Instrumente GmbH & Co. KG, Karlsruhe, Germany)
- Linear actuator: M-229.25S
- Piezo stage: P-545.3D8S + E-727 controller

**Polarization and beam transport**

- BS: 50:50 beamsplitter (Thorlabs Inc., Newton, NJ, USA)
- PBS: polarizing beamsplitter (Thorlabs Inc., Newton, NJ, USA)
- HWP: achromatic half-wave plate (either Thorlabs Inc., Newton, NJ, USA or Edmund Optics Inc., Barrington, NJ, USA)
- QWP: achromatic quarter-wave plate (either Thorlabs Inc., Newton, NJ, USA or Edmund Optics Inc., Barrington, NJ, USA)
- Fibers: polarization maintaining single mode fiber PM-S405-XP and PM780-HP, and multimode fiber M50L022S-A (Thorlabs Inc., Newton, NJ, USA)
- SuK: fiber collimator 60FC-* (Schäfter+Kirchhoff, Hamburg, Germany)

**Lenses and mirrors** (Thorlabs Inc., Newton, NJ, USA if not specified otherwise):
- STG.OBJ: Olympus UPLXAPO 60X (Evident Europe GmbH, Hamburg, Germany)
- L: achromatic lens with VIS or NIR AR coating
- AB.EM: 1-inch elliptical dielectric mirror
- DP.MM: 1-inch metallic mirror
- DP.SQM: 2-inch dielectric square mirrorSTG.EMM: 1-inch elliptical metallic mirror
- mot.: mirror on N-480 mount (Physik Instrumente GmbH & Co. KG, Karlsruhe, Germany)
- E.PP: prism-pair PS875-A

**Dichroic mirrors and filters**
- E.DC: Di03-R561 (Semrock Inc., Rochester, NY, USA)
- E.CU1: clean-up filter LD01-640/8 (Semrock Inc., Rochester, NY, USA)
- E.ND: NE20A (Thorlabs Inc., Newton, NJ, USA)
- AB.DC, BC.DC: Di03-R561 (Semrock Inc., Rochester, NY, USA)
- WFE.DC1: ZT561rdc (Chroma Technology Corp., Bellows Falls, VT, USA)
- WFE.DC2: ZT488rdc (Chroma Technology Corp., Bellows Falls, VT, USA)
- STB.CU1: FBH850-40 (Thorlabs Inc., Newton, NJ, USA)
- STB.CU2: 950-20 bandpass filter (Edmund Optics Inc., Barrington, NJ, USA)
- STB.DC: ZT860lpxr (Chroma Technology Corp., Bellows Falls, VT, USA)
- AC.DC: ZT442rdc (Chroma Technology Corp., Bellows Falls, VT, USA)
- STG.DC1: Di03-R405/488/561/635 (Semrock Inc., Rochester, NY, USA)
- D.NF: quad-notch NF03-405/488/561/635E-25 (Semrock Inc., Rochester, NY, USA)
- D.CU: FESH0800 (Thorlabs Inc., Newton, NJ, USA)
- STG.DC2: T800DCSPXR (Chroma Technology Corp., Bellows Falls, VT, USA)

**Detectors**
- D.APD: SPCM-AQRH-43-BR1 (Excelitas Technologies, Waltham, MA, USA)
- WFD.CAM: DMK 33UX249 (The Imaging Source Europe GmbH, Bremen, Germany)
- STB.CAM: DMK 33UX174 (The Imaging Source Europe GmbH, Bremen, Germany)

**Control hardware and computers** (not shown in the diagram):
- PC: 2 personal computers running Windows 10 (Microsoft Corp., Redmond, WA, USA) and LabVIEW 2020 (National Instruments, Austin, TX, USA)
- DAQ: NI PCIe-6363 (National Instruments, Austin, TX, USA) + USB-3114 (Measurement Computing Corporation, Norton, MA, USA)
- FPGA: NI USB-7856R (National Instruments, Austin, TX, USA)

**Supplementary Table S1.** Focal lengths of lenses in the system.

| Lens | Focal length (mm) |
|---|---|
| E.SuK1, E.SuK2 | 6.2 |
| A.SuK, B.SuK | 7.5 |
| A.L1, B.L1 | 50 |
| A.L2, B.L2 | 30 |
| A.L3, B.L3 | 80 |
| DP.L | 200 |
| OS.L1 | 400 |
| OS.L2 | 200 |
| EOD.L1 | 150 |
| EOD.L2 | 150 |
| EOD.L3 | 200 |
| EOD.L4 | 200 |
| EOD.L5 | 125 |
| EOD.L6 | 400 |
| WFE.SuK1 | 6.2 |
| WFE.L1 | 15 |
| WFE.L2 | 25 |
| WFE.L3 | 100 |
| WFE.L4 | 200 |
| AC.L1 | 4 |
| AC.L2 | 30 |
| AC.L3 | 80 |
| AC.L4 | 200 |
| STG.SL | 180 |
| STG.TL | 300 |
| zS.L | 6.2 |
| STB.L1 | 40 |
| STB.L2 | 6.2 |
| STB.L3 | 150 |
| STB.L4 | 200 |
| STB.L5 | 300 |
| STB.L6 | 80 |
| WFD.L1 | 100 |
| D.L1 | 150 |
| D.L2 | 30 |
| D.L3 | 19 |
| D.L4 | 30 |
| D.L5 | 19 |
| **Waveplate** | **Polarization Axis Orientation (degrees)** |
| A.HWP | ~0 |
| A.QWP | 0 |
| B.QWP | 0 |
| DP.QWP | 45 |
| DP.HWP | 22.5 |
| EOD.HWP | 45 |
| STG.QWP | 45 |

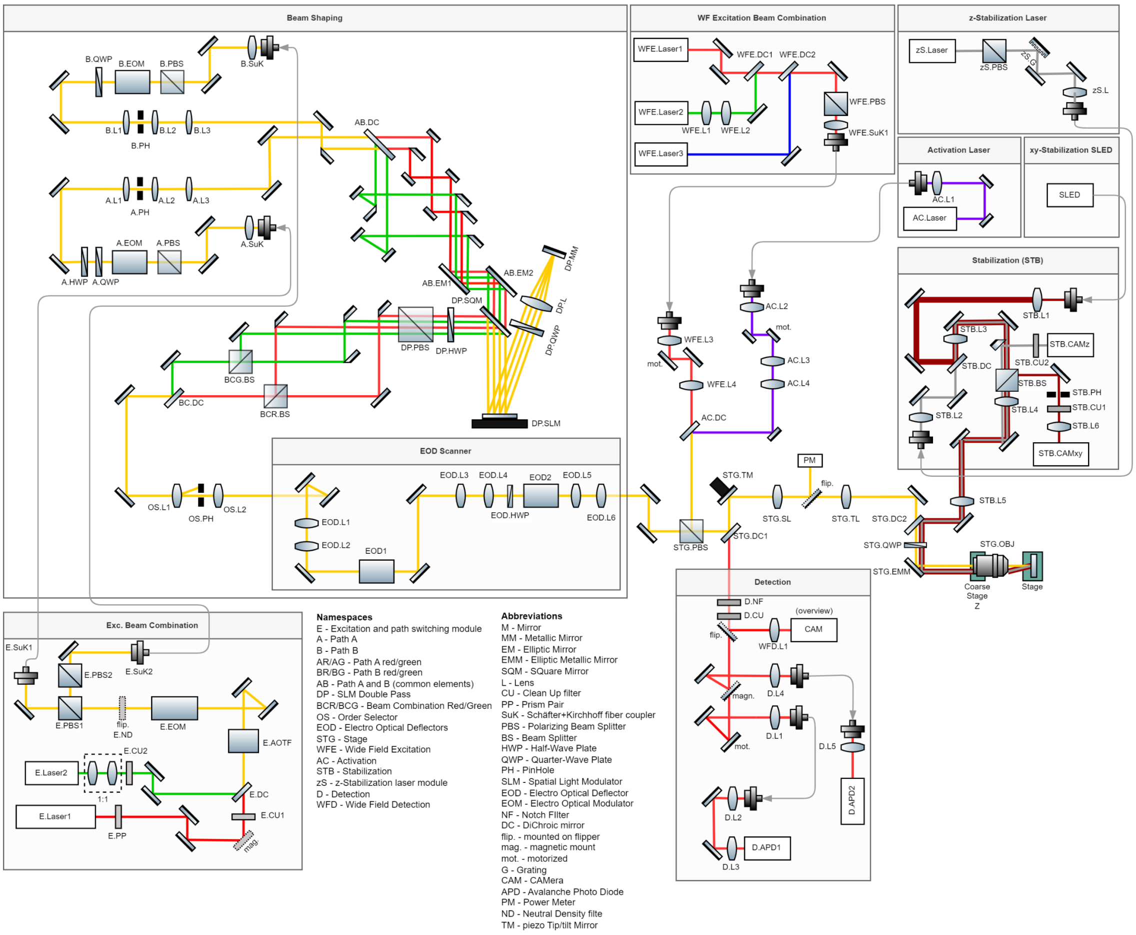

**Supplementary Fig. S13. Detailed optical system.**
The system is arranged into the following functional blocks: Excitation Beam Combination, Beam Shaping, EOD Scanner, Detection, Wide field Excitation Beam Combination, Activation Laser, z-Stabilization Laser, xy-Stabilization SLED.

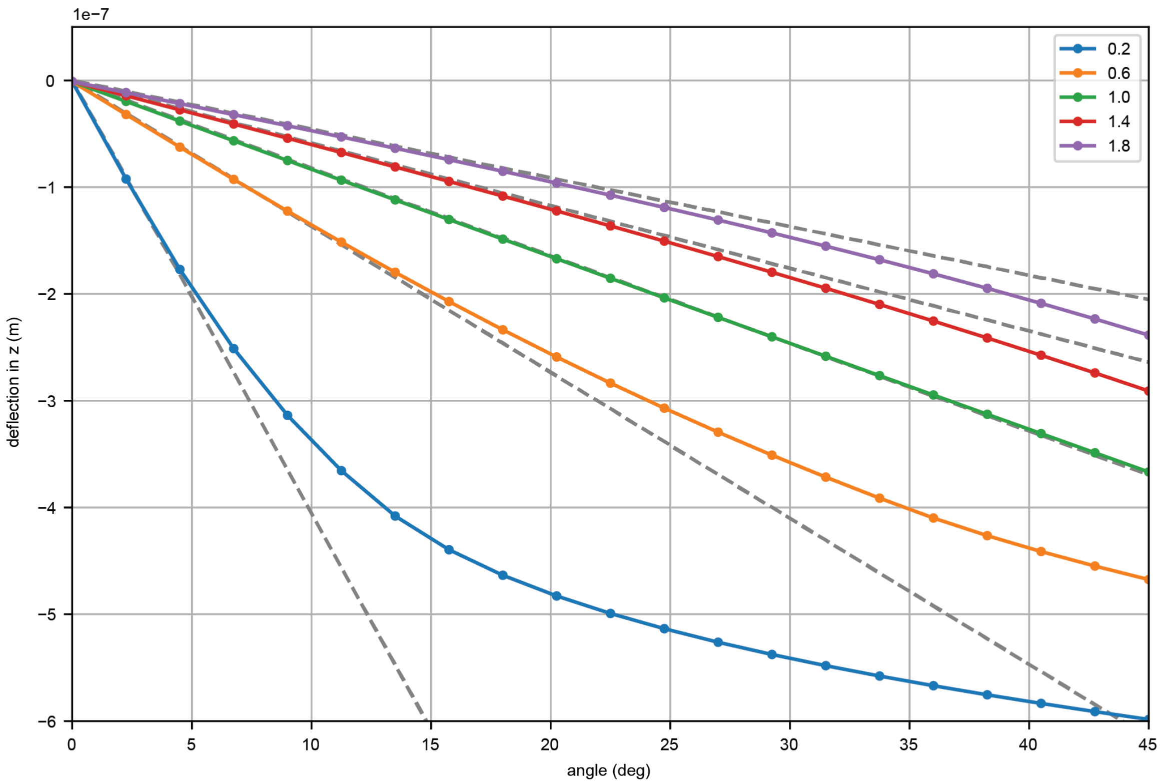

**Supplementary Fig. S14. Simulated nonlinear deflection.**
Deflected distance of the bottle beam zero versus the input polarization's angle. Colored curves correspond to different mixtures of the polarization states at the interference PBS determined by the half-wave plate rotation in front of it. Dashed grey lines are the tangents of the deflection around the origin indicating how each case deviates from linear deflection. When the half-wave plate is at 22.5°, which leads to a relative power of 1 (green line), i.e., an equal mixture of the two polarization states, the deflection is almost perfectly linear. Because of symmetry only positive angles are shown.

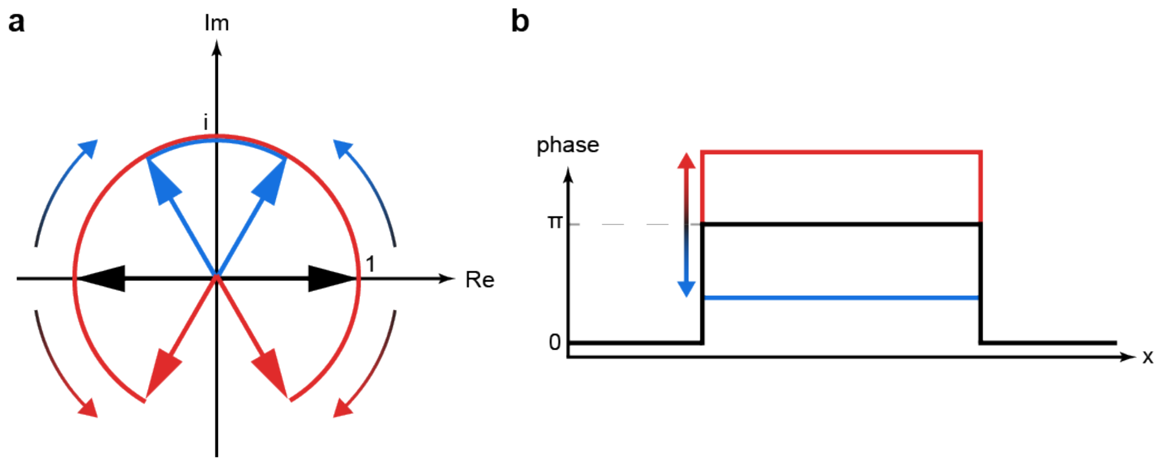

**Supplementary Fig. S15. Bottle beam deflection in the phasor picture.**
**a**, Complex plane with phasors representing the electric field after interference in the two spatial domains $\mathcal{D}_1/\mathcal{D}_2$ of the beam profile. Black corresponds to an undeflected bottle beam; the phase between the phasors is $\pi$. Red and blue correspond to deflected bottle beams (up or down) where the relative phase grows and shrinks, respectively, depending on the input polarization angle. **b**, Profile of the phase distribution of the beam after interference. The $\pi$ phase step grows and shrinks in accordance with the relative phase between phasors shown in a.

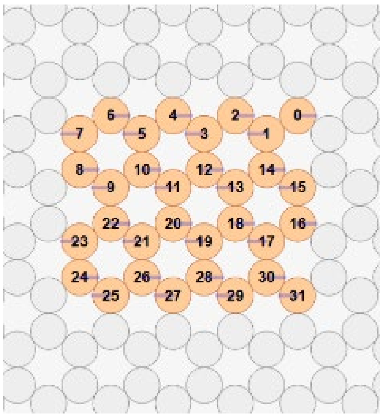

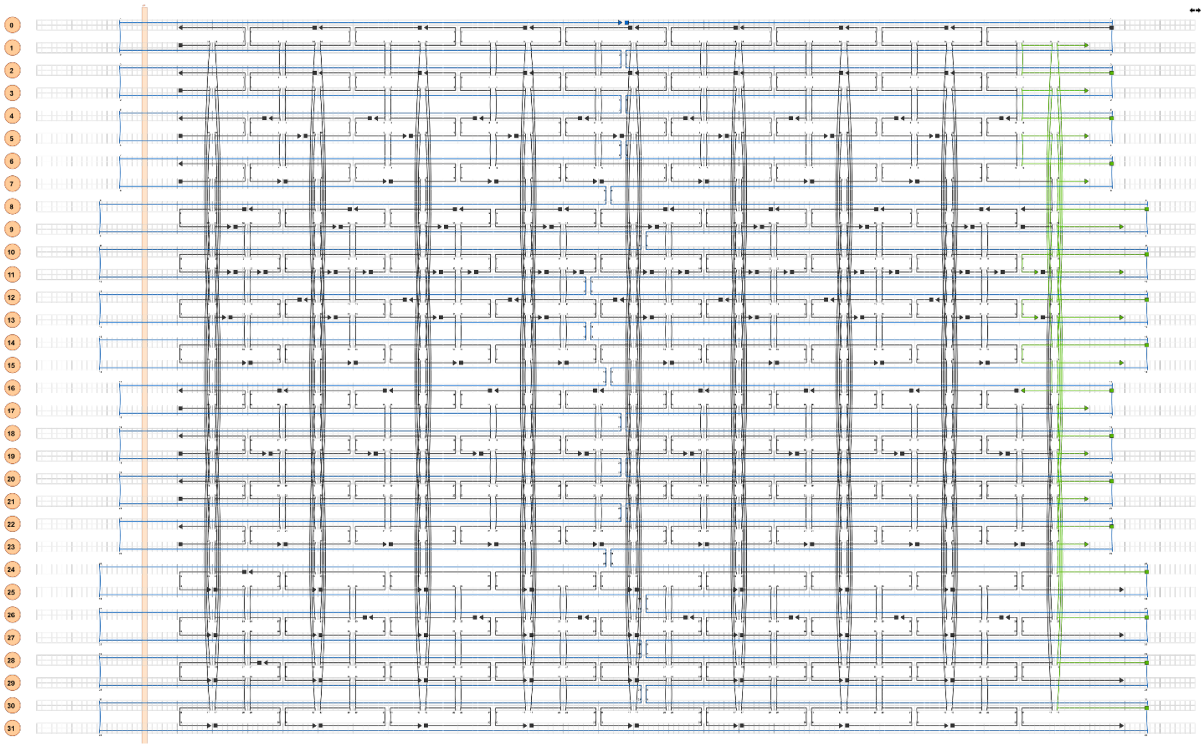

**Supplementary Fig. S16. Cadnano diagram of origami 32HB.**
Biotynilated staples are shown in green.

**Supplementary Table S2.** 2D rectangle origami staple strands.

| Name | Sequence (5’ -> 3’) | Modification |
|---|---|---|
| 23[224]22[240] | GCACAGACAATATTTTTGAATGGGGTCAGTA | |
| 1[128]4[128] | TGACAACTCGCTGAGGCTTGCATTATACCAAGCGCGATGATAAA | |
| 7[96]9[95] | TAAGAGCAAATGTTTAGACTGGATAGGAAGCC | |
| 11[224]13[223] | GCGAACCTCCAAGAACGGGTATGACAATAA | |
| 10[143]9[159] | CCAACAGGAGCGAACCAGACCGGAGCCTTTAC | |
| 11[32]13[31] | AACAGTTTTGTACCAAAAACATTTTATTTC | |
| 10[207]8[208] | ATCCCAATGAGAATTAACTGAACAGTTACCAG | |
| 12[207]10[208] | GTACCGCAATTCTAAGAACGCGAGTATTATTT | |
| 19[160]20[144] | GCAATTCACATATTCCTGATTATCAAAGTGTA | |
| 0[111]1[95] | TAAATGAATTTTCTGTATGGGATTAATTTCTT | |
| 13[64]15[63] | TATATTTTGTCATTGCCTGAGAGTGGAAGATT | |
| 16[207]14[208] | ACCTTTTTATTTTAGTTAATTTCATAGGGCTT | |
| 10[47]8[48] | CTGTAGCTTGACTATTATAGTCAGTTCATTGA | |
| 1[192]4[192] | GCGGATAACCTATTATTCTGAAACAGACGATTGGCCTTGAAGAGCCAC | |
| 18[175]16[176] | CTGAGCAAAAATTAATTACATTTTGGGTTA | |
| 16[47]14[48] | ACAAACGGAAAAGCCCCAAAAACACTGGAGCA | |
| 16[143]15[159] | GCCATCAAGCTCATTTTTTAACCACAAATCCA | |
| 11[256]13[255] | GCCTTAAACCAATCAATAATCGGCACGCGCCT | |
| 8[79]6[80] | AATACTGCCCAAAAGGAATTACGTGGCTCA | |
| 2[79]0[80] | CAGCGAAACTTGCTTTCGAGGTGTTGCTAA | |
| 13[192]15[191] | GTAAAGTAATCGCCATATTTAACAAAACTTTT | |
| 15[256]18[256] | GTGATAAAAAGACGCTGAGAAGAGATAACCTTGCTTCTGTTCGGGAGA | |
| 23[64]22[80] | AAAGCACTAAATCGGAACCCTAATCCAGTT | |
| 15[160]16[144] | ATCGCAAGTATGTAAATGCTGATGATAGGAAC | |
| 6[271]4[272] | ACCGATTGTCGGCATTTTCGGTCATAATCA | |
| 13[256]15[255] | GTTTATCAATATGCGTTATACAAACCGACCGT | |
| 13[32]15[31] | AACGCAAAATCGATGAACGGTACCGGTTGA | |
| 0[143]1[127] | TCTAAAGTTTTGTCGTCTTTCCAGCCGACAA | |
| 10[111]8[112] | TTGCTCCTTTCAAATATCGCGTTTGAGGGGGT | |
| 4[207]2[208] | CCACCCTCTATTCACAAACAAATACCTGCCTA | |
| 20[143]19[159] | AAGCCTGGTACGAGCCGGAAGCATAGATGATG | |
| 6[47]4[48] | TACGTTAAAGTAATCTTGACAAGAACCGAACT | |
| 8[239]6[240] | AAGTAAGCAGACACCACGGAATAATATTGACG | |
| 4[111]2[112] | GACCTGCTCTTTGACCCCCAGCGAGGGAGTTA | |
| 4[143]3[159] | TCATCGCCAACAAAGTACAACGGACGCCAGCA | |
| 3[224]5[223] | TTAAAGCCAGAGCCGCCACCCTCGACAGAA | |
| 21[32]23[31] | TTTTCACTCAAAGGGCGAAAAACCATCACC | |
| 2[143]1[159] | ATATTCGGAACCATCGCCCACGCAGAGAAGGA | |
| 12[47]10[48] | TAAATCGGGATTCCCAATTCTGCGATATAATG | |
| 6[143]5[159] | GATGGTTTGAACGAGTAGTAAATTTACCATTA | |
| 10[271]8[272] | ACGCTAACACCCACAAGAATTGAAAATAGC | |
| 9[96]11[95] | CGAAAGACTTTGATAAGAGGTCATATTTCGCA | |
| 23[128]23[159] | AACGTGGCGAGAAAGGAAGGGAAACCAGTAA | |
| 14[111]12[112] | GAGGGTAGGATTCAAAAGGGTGAGACATCCAA | |
| 2[239]0[240] | GCCCGTATCCGGAATAGGTGTATCAGCCCAAT | |
| 5[32]7[31] | CATCAAGTAAAACGAACTAACGAGTTGAGA | |
| 17[160]18[144] | AGAAAACAAAGAAGATGATGAAACAGGCTGCG | |
| 20[271]18[272] | CTCGTATTAGAAATTGCGTAGATACAGTAC | |
| 4[255]6[248]-biotin | AGCCACCACTGTAGCGCGTTTTCAAGGGAGGGAAGGTAAA | 5’-biotin |
| 17[96]19[95] | GCTTTCCGATTACGCCAGCTGGCGGCTGTTTC | |
| 19[96]21[95] | CTGTGTGATTGCGTTGCGCTCACTAGAGTTGC | |
| 3[32]5[31] | AATACGTTTGAAAGAGGACAGACTGACCTT | |
| 4[79]2[80] | GCGCAGACAAGAGGCAAAAGAATCCCTCAG | |
| 1[32]3[31] | AGGCTCCAGAGGCTTTGAGGACACGGGTAA | |
| 1[256]4[256] | CAGGAGGTGGGGTCAGTGCCTTGAGTCTCTGAATTTACCGGGAACCAG | |
| 18[255]20[248]-biotin | AACAATAACGTAAAACAGAAATAAAAATCCTTTGCCCGAA | 5’-biotin |
| 18[191]20[184]-biotin | ATTCATTTTTGTTTGGATTATACTAAGAAACCACCAGAAG | 5’-biotin |
| 16[175]14[176] | TATAACTAACAAAGAACGCGAGAACGCCAA | |
| 1[64]4[64] | TTTATCAGGACAGCATCGGAACGACACCAACCTAAAACGAGGTCAATC | |
| 6[207]4[208] | TCACCGACGCACCGTAATCAGTAGCAGAACCG | |
| 20[207]18[208] | GCGGAACATCTGAATAATGGAAGGTACAAAAT | |
| 23[192]22[208] | ACCCTTCTGACCTGAAAGCGTAAGACGCTGAG | |
| 3[96]5[95] | ACACTCATCCATGTTACTTAGCCGAAAGCTGC | |
| 21[160]22[144] | TCAATATCGAACCTCAAATATCAATTCCGAAA | |
| 21[248]23[255] | AGATTAGAGCCGTCAAAAAACAGAGGTGAGGCCTATTAGT | |
| 23[256]22[272] | CTTTAATGCGCGAACTGATAGCCCCACCAG | |

| | | |
|---|---|---|
| 22[111]20[112] | GCCCGAGAGTCCACGCTGGTTTGCAGCTAACT | |
| 2[47]0[48] | ACGGCTACAAAAGGAGCCTTTAATGTGAGAAT | |
| 21[184]23[191] | TCAACAGTTGAAAGGAGCAAATGAAAAATCTAGAGATAGA | |
| 22[175]20[176] | ACCTTGCTTGGTCAGTTGGCAAAGAGCGGA | |
| 18[47]16[48] | CCAGGGTTGCCAGTTTGAGGGGACCCGTGGGA | |
| 12[143]11[159] | TTCTACTACGCGAGCTGAAAAGGTTACCGCGC | |
| 16[79]14[80] | GCGAGTAAAAATATTTAAATTGTTACAAAG | |
| 17[224]19[223] | CATAAATCTTTGAATACCAAGTGTTAGAAC | |
| 12[271]10[272] | TGTAGAAATCAAGATTAGTTGCTCTTACCA | |
| 10[175]8[176] | TTAACGTCTAACATAAAAACAGGTAACGGA | |
| 2[111]0[112] | AAGGCCGCTGATACCGATAGTTGCGACGTTAG | |
| 8[111]6[112] | AATAGTAAACACTATCATAACCCTCATTGTGA | |
| 8[271]6[272] | AATAGCTATCAATAGAAAATTCAACATTCA | |
| 7[120]9[127] | CGTTTACCAGACGACAAAGAAGTTTTGCCATAATTCGA | |
| 7[160]8[144] | TTATTACGAAGAACTGGCATGATTGCGAGAGG | |
| 18[271]16[272] | CTTTTACAAAATCGTCGCTATTAGCGATAG | |
| 22[207]20[208] | AGCCAGCAATTGAGGAAGGTTATCATCATTTT | |
| 0[175]0[144] | TCCACAGACAGCCCTCATAGTTAGCGTAACGA | |
| 22[47]20[48] | CTCCAACGCAGTGAGACGGGCAACCAGCTGCA | |
| 0[207]1[191] | TCACCAGTACAAACTACAACGCCTAGTACCAG | |
| 3[160]4[144] | TTGACAGGCCACCACCAGAGCCGCGATTTGTA | |
| 13[128]15[127] | GAGACAGCTAGCTGATAAATTAATTTTTGT | |
| 14[143]13[159] | CAACCGTTTCAAATCACCATCAATTCGAGCCA | |
| 15[128]18[127] | TAAATCAAAATAATTCGCGTCTCGGAAACCAGGCAAAGGGAAGGG | |
| 5[160]6[144] | GCAAGGCCTCACCAGTAGCACCATGGGCTTGA | |
| 9[64]11[63] | CGGATTGCAGAGCTTAATTGCTGAAACGAGTA | |
| 2[271]0[272] | GTTTTAACTTAGTACCGCCACCCAGAGCCA | |
| 7[184]9[191] | CGTAGAAAATACATACCGAGGAAACGCAATAAGAAGCGCA | |
| 16[239]14[240] | GAATTTATTTAATGGTTTGAAATATTCTTACC | |
| 18[111]16[112] | TCTTCGCTGCACCGCTTCTGGTGCGGCCTTCC | |
| 0[271]1[255] | CCACCCTCATTTTCAGGGATAGCAACCGTACT | |
| 0[47]1[31] | AGAAAGGAACAACTAAAGGAATTCAAAAAAA | |
| 1[224]3[223] | GTATAGCAAACAGTTAATGCCCAATCCTCA | |
| 9[160]10[144] | AGAGAGAAAAAAATGAAAATAGCAAGCAAACT | |
| 21[56]23[63] | AGCTGATTGCCCTTCAGAGTCCACTATTAAAGGGTGCCGT | |
| 22[239]20[240] | TTAACACCAGCACTAACAACTAATCGTTATTA | |
| 7[248]9[255] | GTTTATTTTGTCACAATCTTACCGAAGCCCTTTAATATCA | |
| 10[239]8[240] | GCCAGTTAGAGGGTAATTGAGCGCTTTAAGAA | |
| 13[224]15[223] | ACAACATGCCAACGCTCAACAGTCTTCTGA | |
| 2[175]0[176] | TATTAAGAAGCGGGGTTTTGCTCGTAGCAT | |
| 12[111]10[112] | TAAATCATATAACCTGTTTAGCTAACCTTTAA | |
| 22[79]20[80] | TGGAACAACCGCCTGGCCCTGAGGCCCGCT | |
| 0[239]1[223] | AGGAACCCATGTACCGTAACACTTGATATAA | |
| 12[239]10[240] | CTTATCATTCCCGACTTGCGGGAGCCTAATTT | |
| 11[192]13[191] | TATCCGGTCTCATCGAGAACAAGCGACAAAAG | |
| 18[126]20[120]-biotin | CGATCGGCAATTCCACACAACAGGTGCCTAATGAGTG | 5'-biotin |
| 17[32]19[31] | TGCATCTTTCCCAGTCACGACGGCCTGCAG | |
| 20[239]18[240] | ATTTTAAAATCAAAATTATTTGCACGGATTCG | |
| 4[271]2[272] | AAATCACCTTCCAGTAAGCGTCAGTAATAA | |
| 18[207]16[208] | CGCGCAGATTACCTTTTTTAATGGGAGAGACT | |
| 8[143]7[159] | CTTTTGCAGATAAAAACCAAAATAAAGACTCC | |
| 11[160]12[144] | CCAATAGCTCATCGTAGGAATCATGGCATCAA | |
| 14[47]12[48] | AACAAGAGGGATAAAAATTTTTAGCATAAAGC | |
| 15[96]17[95] | ATATTTTGGCTTTCATCAACATTATCCAGCCA | |
| 15[224]17[223] | CCTAAATCAAAATCATAGGTCTAAACAGTA | |
| 16[111]14[112] | TGTAGCCATTAAAATTCGCATTAAATGCCGGA | |
| 4[239]2[240] | GCCTCCCTCAGAATGGAAAGCGCAGTAACAGT | |
| 6[175]4[176] | CAGCAAAAGGAAACGTCACCAATGAGCCGC | |
| 8[47]6[48] | ATCCCCCTATACCACATTCAACTAGAAAAATC | |
| 14[175]12[176] | CATGTAATAGAATATAAAGTACCAAGCCGT | |
| 6[111]4[112] | ATTACCTTTGAATAAGGCTTGCCCAAATCCGC | |
| 4[63]6[56]-biotin | ATAAGGGAACCGGATATTCATTACGTCAGGACGTTGGGAA | 5'-biotin |
| 20[79]18[80] | TTCCAGTCGTAATCATGGTCATAAAAGGGG | |
| 9[192]11[191] | TTAGACGGCCAAATAAGAAACGATAGAAGGCT | |
| 19[224]21[223] | CTACCATAGTTTGAGTAACATTTAAAATAT | |
| 14[271]12[272] | TTAGTATCACAATAGATAAGTCCACGAGCA | |
| 18[63]20[56]-biotin | ATTAAGTTTACCGAGCTCGAATTCGGGAAACCTGTCGTGC | 5'-biotin |
| 20[111]18[112] | CACATTAAAATTGTTATCCGCTCATGCGGGCC | |
| 12[175]10[176] | TTTTATTTAAGCAAATCAGATATTTTTTGT | |
| 8[207]6[208] | AAGGAAACATAAAGGTGGCAACATTATCACCG | |

| | | |
|---|---|---|
| 2[207]0[208] | TTTCGGAAGTGCCGTCGAGAGGGTGAGTTTCG | |
| 9[128]11[127] | GCTTCAATCAGGATTAGAGAGTTATTTTCA | |
| 7[224]9[223] | AACGCAAAGATAGCCGAACAAACCCTGAAC | |
| 1[160]2[144] | TTAGGATTGGCTGAGACTCCTCAATAACCGAT | |
| 20[175]18[176] | ATTATCATTCAATATAATCCTGACAATTAC | |
| 14[239]12[240] | AGTATAAAGTTCAGCTAATGCAGATGTCTTTC | |
| 19[32]21[31] | GTCGACTTCGGCCAACGCGCGGGGTTTTTC | |
| 4[47]2[48] | GACCAACTAATGCCACTACGAAGGGGGTAGCA | |
| 15[192]18[192] | TCAAATATAACCTCCGGCTTAGGTAACAATTTCATTTGAAGGCGAATT | |
| 6[239]4[240] | GAAATTATTGCCTTTAGCGTCAGACCGGAACC | |
| 0[79]1[63] | ACAACTTTCAACAGTTTCAGCGGATGTATCGG | |
| 4[191]6[184]-biotin | CACCCTCAGAAACCATCGATAGCATTGAGCCATTTGGGAA | 5’-biotin |
| 22[143]21[159] | TCGGCAAATCCTGTTTGATGGTGGACCCTCAA | |
| 7[56]9[63] | ATGCAGATACATAACGGGAATCGTCATAAATAAAGCAAAG | |
| 15[32]17[31] | TAATCAGCGGATTGACCGTAATCGTAACCG | |
| 23[160]22[176] | TAAAAGGGACATTCTGGCCAACAAAGCATC | |
| 1[96]3[95] | AAACAGCTTTTTGCGGGATCGTCAACACTAAA | |
| 12[79]10[80] | AAATTAAGTTGACCATTAGATACTTTTGCG | |
| 23[32]22[48] | CAAATCAAGTTTTTTGGGGTCGAAACGTGGA | |
| 13[160]14[144] | GTAATAAGTTAGGCAGAGGCATTTATGATATT | |
| 21[120]23[127] | CCCAGCAGGCGAAAAATCCCTTATAAATCAAGCCGGCG | |
| 21[224]23[223] | CTTTAGGGCCTGCAACAGTGCCAATACGTG | |
| 23[96]22[112] | CCCGATTTAGAGCTTGACGGGGAAAAAGAATA | |
| 11[128]13[127] | TTTGGGGATAGTAGTAGCATTAAAAGGCCG | |
| 20[47]18[48] | TTAATGAACTAGAGGATCCCCGGGGGGTAACG | |
| 5[96]7[95] | TCATTCAGATGCGATTTTAAGAACAGGCATAG | |
| 11[96]13[95] | AATGGTCAACAGGCAAGGCAAAGAGTAATGTG | |
| 13[96]15[95] | TAGGTAAACTATTTTTGAGAGATCAAACGTTA | |
| 4[175]2[176] | CACCAGAAAGGTTGAGGCAGGTCATGAAAG | |
| 15[64]18[64] | GTATAAGCCAACCCGTCGGATTCTGACGACAGTATCGGCCGCAAGGCG | |
| 9[224]11[223] | AAAGTCACAAAATAAACAGCCAGCGTTTTA | |
| 18[143]17[159] | CAACTGTTGCGCCATTCGCCATTCAAACATCA | |
| 9[256]11[255] | GAGAGATAGAGCGTCTTTCCAGAGGTTTTGAA | |
| 18[79]16[80] | GATGTGCTTCAGGAAGATCGCACAATGTGA | |
| 14[79]12[80] | GCTATCAGAAATGCAATGCCTGAATTAGCA | |
| 10[79]8[80] | GATGGCTTATCAAAAAGATTAAGAGCGTCC | |
| 4[127]6[120]-biotin | TTGTGTCGTGACGAGAAACACCAAATTTCAACTTTAAT | 5’-biotin |
| 7[32]9[31] | TTTAGGACAAATGCTTTAAACAATCAGGTC | |
| 8[175]6[176] | ATACCCAACAGTATGTTAGCAAATTAGAGC | |
| 11[64]13[63] | GATTTAGTCAATAAAGCCTCAGAGAACCCTCA | |
| 14[207]12[208] | AATTGAGAATTCTGTCCAGACGACTAAACCAA | |
| 18[239]16[240] | CCTGATTGCAATATATGTGAGTGATCAATAGT | |
| 16[271]14[272] | CTTAGATTTAAGGCGTTAAATAAAGCCTGT | |
| 6[79]4[80] | TTATACCACCAAATCAACGTAACGAACGAG | |
| 9[32]11[31] | TTTACCCCAACATGTTTTAAATTTCCATAT | |
| 5[224]7[223] | TCAAGTTTCATTAAAGGTGAATATAAAAGA | |
| 21[96]23[95] | AGCAAGCGTAGGGTTGAGTGTTGTAGGGAGCC | |
| 22[271]20[272] | CAGAAGATTAGATAATACATTTGTCGACAA | |

**Supplementary Table S3.** 3D 32 HB origami staple strands.

| Name | Sequence (5' -> 3') | Modification |
|---|---|---|
| 29[35]15[41]-32HB | GGAGGTTCCCTCATAAATGAAATTTATCCGAACCT | |
| 4[66]7[69]-32HB | ATTGACCAAGTAACAATTTTCCCTT | |
| 13[38]0[28]-32HB | GCATCAAATCACTAAAATGGTCAGTTGGCAAA | |
| 8[104]5[94]-32HB | TAGTGAAAGATTAAAAACAAACATCAAGATTA | |
| 13[59]0[56]-32HB | ATCGATTACCGTAACAACTCAAACC | |
| 19[28]29[34]-32HB | AGCCCTTAGGGTTGGTACTCA | |
| 31[161]26[172]-32HB | GGCTCCAAAAGGAGCCCACGCAGGGTAGCCTAAAAGCCTGATAAG | |
| 23[49]25[55]-32HB | TAGCAGCAATCACCCCTCAGA | |
| 12[178]31[181]-32HB | AACTGGTAGCTGGTCATTATTCTGCACCATCGCCTTTAA | |
| 26[192]9[185]-32HB | CGAAGAATAAGTTTAATTTCAACTTGATTTAGCATAGTACAGT | |
| 2[76]16[70]-32HB | CGAACGTTTAGAGCTCATCGTAAATCAACCTAATT | |
| 25[119]11[121]-32HB | CGCAAGATTCATATCAAAAGAATTGCATTTA | |
| 28[221]19[209]-32HB-biotin | TTTCCATTAAACGGGTAAACAGGTCA | 5'-biotin |
| 23[154]25[160]-32HB | ATTACAGTGCGATTTACCCAA | |
| 14[221]1[209]-32HB-biotin | GGGAGAAGCCTTTATTTCAAGTTTTT | 5'-biotin |
| 8[167]5[157]-32HB | GCGCAACGGTTTTCATCATGGTCATAGCTGGG | |
| 11[45]8[42]-32HB | CCAACAAATTCACTTTTTGGTTGGG | |
| 16[69]19[66]-32HB | TGCCAGTAATAACACAAGAATAAAG | |
| 13[185]0[182]-32HB | TTTGATTTTTACGTGAACGAACCCT | |
| 28[44]24[42]-32HB | TGCGTCAGTGCGTTCCACCACCACGGAACCA | |
| 29[119]15[125]-32HB | AGCTTTTGGCCGCCTATGCAAATTAACACATACAG | |
| 2[55]16[49]-32HB | GTTTGAGGGAGCACCGCCCAATGCGGGATAAACAG | |
| 21[28]27[34]-32HB | TTGTCACTCATACATAAGTTT | |
| 16[174]19[171]-32HB | TTGGGGCATTCCCATTTGCGGGCTT | |
| 23[133]25[139]-32HB | ACGAACTATTATACTGACAAG | |
| 25[161]11[163]-32HB | ATCAACGCAGATGAACTGGATATCAGGTTTT | |
| 27[35]13[37]-32HB | TAACGGGCGTCGAGTTTAAGAAGCAAGACGA | |
| 0[139]4[130]-32HB | AACGTGGCGAGAAAACGTGGATGGAACAGGGCAACTGG | |
| 23[196]25[216]-32HB | ACATTCAGAGATGGGCTTGCCCTGACGAGAAACAC | |
| 5[137]29[139]-32HB | GCAATTGACCTTAATATCAGAAGCGCCCGAAACTCATCCGTCACC | |
| 17[28]31[34]-32HB | ACGTCAAAGTTAGCCGTTAGT | |
| 4[192]8[196]-32HB | GCTGTGCGTTGATCCCCGGACGGCCAGTGCCAGGAAACC | |
| 31[35]28[45]-32HB | AAATGAATTTTCTGCAGACAGTAGTACCTAAG | |
| 23[28]25[34]-32HB | CGTCACCCACCGGACGCCACC | |
| 19[151]23[153]-32HB | ATCCTGACTAAGTAAAACCAGACGCAACATT | |
| 27[140]13[142]-32HB | ACAAAGTCTAAAACAGACTTCGAATATATCA | |
| 11[200]29[216]-32HB | CCAACAGTTCACTCCAAATACGTAGGACTAAAGACTTTTTCATGA | |
| 24[221]23[209]-32HB-biotin | CGAGTAGTAAATTGGGCTTACTAATG | 5'-biotin |
| 0[160]4[151]-32HB | GCTTGACGGGGAAAGGCGAAAGGGTTGACACCGCCTTT | |
| 9[186]12[179]-32HB | ATCATGTGAGCATTAATCATTAATGAATCGGAAGCGGTTAA | |
| 4[45]7[48]-32HB | AACGATTCGCCTTTTTTAAATCGTC | |
| 5[179]29[181]-32HB | TCACGAGTAAGCTCATTACCATAAAGCGAACGCACCAACAACGGC | |
| 9[144]12[137]-32HB | ATCACGGCGGTAAAGTGATTGGGCGCCAGGGAGCTGATGAA | |
| 2[214]16[196]-32HB-biotin | GTTTGATGGTGGTTCCGAACAAATCAACGCAAGACATTATTCAATAA | 5'-biotin |
| 27[119]13[121]-32HB | TGCGATTCCCAGAAAAGATTACATGTTTAAA | |
| 0[55]4[46]-32HB | CTCAATCAATATCTATCTTTATAACATTTCCTGATTTT | |
| 25[35]11[38]-32HB | CTCAGAGGTAAGCGAATCAATATAAAAGTTTA | |
| 19[88]23[90]-32HB | AACCGCAGTAACATTCAAATTATCTTGCCTT | |
| 13[101]0[98]-32HB | CAAGATCGAGAAGGATTTCAAATGA | |
| 5[28]20[28]-32HB | AGAAACATATAAAGCGCCATAAAACGCA | |
| 0[118]4[109]-32HB | AAGAGAGCCAGCAGAGAAGTACAAACAAATAATGGACC | |
| 7[91]27[97]-32HB | ATAGCTTTTTATCATGGTTTGAAAGGTGACCGATTAATAAATCTATTTC | |
| 13[122]0[119]-32HB | TCACAGGGTGATTAAAGAGGAAGGG | |
| 7[175]27[181]-32HB | CGACGTTTCGCCATACGACGAAGAGCAAAATACTGACTTTGAAAATTGT | |
| 11[185]2[182]-32HB | TAACGTAATCGCCACGCTAAATCCC | |
| 31[119]26[130]-32HB | TGCGAATAATAATTAGTTAAAGCGGGATTTTGACCATACCAAGCT | |
| 29[161]15[167]-32HB | CGGAACGATAACCGACAGTTGGCGAGCTCCTCAGA | |
| 12[157]31[160]-32HB | TACCTGATAAAAGAGCTTCCATATAATATATTAAAAAAA | |
| 15[42]11[44]-32HB | CCCGACTTAGCAAGGTAGAAAGCCTGTTACG | |
| 26[171]9[164]-32HB | AGGATAACAAATGTGAATTACCTTAGTAGAAAATAACCCGCCA | |
| 11[66]8[63]-32HB | GGCATTTAGTATTAATTTACCTTTT | |
| 11[122]2[119]-32HB | AATTACAGGAAGTGAGACAGAGTCC | |
| 19[67]23[69]-32HB | TTAGAAAATAAGCGCCAGAGCCATTCAGTAG | |
| 13[80]0[77]-32HB | TTCCTTTATTTCGTCAATCACCTTG | |
| 20[214]11[216]-32HB-biotin | CAAATGCTTTAATCAAAAATAATTCG | 5'-biotin |
| 27[98]13[100]-32HB | GGAACCTGAGACTCTAACGGAGGGTAATAAC | |

| | | |
|---|---|---|
| 29[56]15[62]-32HB | TCAGAACCCTGTAGCAGAGAGTACAAAAGGTTTTG | |
| 15[84]11[86]-32HB | TGCTATTAGCCGTTAAGAACGAGACGACATA | |
| 31[77]26[88]-32HB | AGTTTCAGCGGAGTAGTTTCGCCTCAGAGAAGGATCCCCTGCCCT | |
| 8[221]7[209]-32HB-biotin | CGGCACCGCTTCTGGTGCCAGCTTGC | 5’-biotin |
| 11[101]2[98]-32HB | ACCGAAGTAATCTTCTGATTCGACA | |
| 15[147]11[149]-32HB | ATTAAGCCTGAGTATTCTAGCCCGGTTGAAA | |
| 25[98]11[100]-32HB | AGGAGGTACAAACAGAGGGAGAAGACTCAGT | |
| 7[28]22[28]-32HB | ACCTTGCACTATATAAGAACGGTAGCAC | |
| 15[189]11[191]-32HB | ACCAAAAGATAAAAAGAGATCATGAACGCAA | |
| 9[39]4[28]-32HB | AAATTACCAGATAACGGTCAGATGAATATAC | |
| 9[165]12[158]-32HB | GTTTCGGATTCCTAATGGCGGGGAGAGGCGGTGGCCCTATG | |
| 8[125]27[118]-32HB | ATTACGCCAGCTAATGATAAATTTTGACAAATATGTGGCCTTTGAAACA | |
| 26[87]9[80]-32HB | CATTCCGCCAGCCCCTTATTAGCGTATCAAGTACCGTCATGAC | |
| 7[70]27[76]-32HB | AGAATCCAGAGACTCATCTTCCCGACTTAAGACAAGAATGGATAAACAG | |
| 11[143]2[140]-32HB | GTTAATAATCATGCCCTTGTGTTGT | |
| 7[154]27[160]-32HB | ACGCCAGTGTTGGGTGCATCTTCGTTTATGTTTAGACGGTGTAGATTTG | |
| 4[171]7[174]-32HB | ACGCAGTGAGCATTCGTACCAGTCA | |
| 2[139]16[133]-32HB | TCCAGTTCTCCAACGGTAAAGAAAGAATCTAATAG | |
| 23[175]25[181]-32HB | CAGTTGATAATCATGCTGCTC | |
| 30[221]17[209]-32HB-biotin | AAACAGCTTGATACCGATACCATTAG | 5’-biotin |
| 9[123]12[116]-32HB | CACGATAGAGACAACATTCTTTTCACTTAGAAAGGGCAGAT | |
| 0[76]4[67]-32HB | CTGAACCTCAAATATAATAGATATTAATAATTCATGAA | |
| 29[182]15[188]-32HB | TACAGAGGACAACAGAACGAGAGCTATACGGTTGT | |
| 4[129]7[132]-32HB | TTTTACGAGCCATCCGCTATGTGCT | |
| 8[41]6[28]-32HB | TTATATATTCTGTAATGGAAACAGTACA | |
| 27[77]13[79]-32HB | TTAATGCTAGGATTGAAACCGAGAGAGATCA | |
| 11[164]2[161]-32HB | TTGTCAATCATGAGAGAGAATAGCC | |
| 26[65]9[59]-32HB | GCACCAGAACCTTTTCATAATCAAACCGTAATTGGGAAATTT | |
| 29[98]15[104]-32HB | CCCTCATGTACCGTGAATTAAACCAACGCAGCTAC | |
| 31[140]26[151]-32HB | GTTGAAAATCTCCACGGTCGCGCGAAAGGAATACAACAACGGACA | |
| 12[52]31[55]-32HB | ACGCCCAATCAAGCCCAAGCCTTTACATTCCATATGGGA | |
| 2[97]16[91]-32HB | ACTCGTAACATTTGACAAGCATTGCACCCTAACGA | |
| 16[153]19[150]-32HB | TGGCATCTTTCATTAATTGCTAAAT | |
| 31[182]26[193]-32HB | TTGTATCGGTTTATCGACAATGCTTTGAATGCCACTCCGCGAAAC | |
| 12[73]31[76]-32HB | AACATTCCTTATAACCCATAAAAACTACAAACTTTCAAC | |
| 0[181]4[172]-32HB | AAAGGGAGCCCCCGGCGATGGTCAAAAGTTGCAGCCCA | |
| 19[46]23[48]-32HB | TAAGCAACATAGAAAATCAGCAAACCATCGA | |
| 26[221]21[209]-32HB-biotin | GAGGCGCAGACGGTCAATCTTGAATC | 5’-biotin |
| 15[168]11[170]-32HB | GCATAAACATATATCGGAGAGAGCATGTTAA | |
| 25[56]11[58]-32HB | GCCGCCAGTCTCTGGTTTACCCATACATTTT | |
| 23[91]25[97]-32HB | TAGCGTCTCATAGCCATTGAC | |
| 25[140]11[142]-32HB | AACCGGAAGGCGCAGGGTAATTTATAGTTTT | |
| 9[60]12[53]-32HB | TAGTCATATGTTTGAATCGTAGATTTTCAGGTATCAGAAGA | |
| 8[83]5[73]-32HB | AGGTCTGTTGAAAAAATTAATTACATTTTACA | |
| 16[195]19[192]-32HB | CCTGTTTTAGATTTTCCTTTTGAAG | |
| 4[87]7[90]-32HB | ACGTGAGGCGAAAAACAACATAGCG | |
| 12[115]31[118]-32HB | TGTAGACAGTCTAAACAACCCTGAACAATAGGAAGGAAT | |
| 23[70]25[76]-32HB | CGACAGATTGCCATCACCACC | |
| 11[80]2[77]-32HB | AGTAGACAATAATCCTGACTTTGCC | |
| 18[214]13[216]-32HB-biotin | AGAGAGTACCTTGCTATCAGGTCATT | 5’-biotin |
| 29[77]15[83]-32HB | CCGCCACTCACCAGAGGGAAGTCCAGAGGATTAGT | |
| 9[81]12[74]-32HB | CTACTAGAAATCGCGCAAAAACAGAAATAAACAATATAAAC | |
| 11[192]8[189]-32HB | TAGGACATTAAGGCCTCAAGGCAAA | |
| 4[214]13[199]-32HB-biotin | AGTCGGGAAACCTGTCGTGCCCAGCAAGAATCGTACA | 5’-biotin |
| 11[59]2[56]-32HB | AGGCCTAATGCTGATGGCTTTAAAA | |
| 5[74]29[76]-32HB | AAAAAGCCTGTTTTCGAAAACGTACCAGAAGAGCGGGGCTCAGAA | |
| 5[158]29[160]-32HB | GTGCTCCGTGCATTAAACTTTACCGCGTTTTGGCAAAAACAGCAT | |
| 13[200]31[216]-32HB | AAGTAATTGCAGTTTGAGTTGCGCCAGCTTGCTTTCGAGGTGAAT | |
| 9[196]27[216]-32HB | GGAAGATAAGGAATATATTCAATAAGGGCCTGCTCCATGTTACTTAGCC | |
| 13[164]0[161]-32HB | ATGCTTTAAATTATCAGGATTTAGA | |
| 4[108]7[111]-32HB | TACCTTACCTGGATGATGGACGCTG | |
| 27[56]13[58]-32HB | TAACAGTCAGTACCGCCGAACTGAGTTAATA | |
| 25[182]11[184]-32HB | ATTCAGTCTGACCACGGAATCGAGAATGTTT | |
| 11[39]2[28]-32HB | ACATATCAACATCATATATCATTTTGCGGAA | |
| 26[129]9[122]-32HB | GACCGTAATCTCAGTCAGGACGTTGTCTACGTAGAGGCTTAGT | |
| 27[161]13[163]-32HB | TATCATCCGAAAGAAATTCGAATGGCTTTTA | |
| 7[133]27[139]-32HB | GCAAGGCCGGGCCTTAGATGGAAACCAACCAGAGGTAGGCTGGCGCGAA | |
| 19[193]23[195]-32HB | CAAAGAAAACGTCATAATACGAGGGAATACC | |
| 26[150]9[143]-32HB | GACCTATTCATTTAAGAACTGGCTCAACGGAAACGATAAGCGC | |
| 2[181]16[175]-32HB | TTATAAACCCACTAGAACCCTGCTAAATTTTTCAT | |

| | | |
|---|---|---|
| 16[111]19[108]-32HB | GAATCTTCTGAACAAGTCAGAATAC | |
| 12[136]31[139]-32HB | AAGCATGATATATGCTGTACGGTGTTGCAGGGTTTTCAC | |
| 1[28]16[28]-32HB | AGGTTATGATATAGGTTTTAGCCAATCC | |
| 16[48]19[45]-32HB | CCATATTAATAGCATAATAAGAAAG | |
| 7[112]5[115]-32HB | AGGGCGAAAGGGGGCACAATTCCAAGAAAGCA | |
| 19[130]23[132]-32HB | GAAAAAGCGGAGTTTTGAATAGCGTAATAAA | |
| 13[143]0[140]-32HB | ACCGATGTGTAGTCAAAGGCCGGCG | |
| 6[214]11[199]-32HB-biotin | TGCAGGTCGACTCTAGAGGCGCTCACTTCATCAAACG | 5’-biotin |
| 11[150]8[147]-32HB | TTCGGGAACAAGTAACCGAAGGGCG | |
| 9[102]12[95]-32HB | ACCTAAACACATTTCAAATATCAAAATTATTGATTATATCT | |
| 12[221]3[209]-32HB-biotin | AGAGTCTGGAGCAAACAAGGGCGAAA | 5’-biotin |
| 8[62]5[52]-32HB | TAACCTCATTAATTTCATTTGAATTACCTGAT | |
| 10[221]5[209]-32HB-biotin | GGCCTTCCTGTAGCCAGCTTGCCCGC | 5’-biotin |
| 16[214]15[216]-32HB-biotin | TTTCGCAAATGGGACCCTGTAATACT | 5’-biotin |
| 7[49]27[55]-32HB | GCTATTACGGCTTACAAATATTTAGAGCTCATATGAATTTACCCTTGAG | |
| 0[97]4[88]-32HB | AAAATCTAAAGCATAGATAATTTAAATCTTGTTTGTGC | |
| 11[129]8[126]-32HB | AACGGTAATGGGTTGGTGCTTCGCT | |
| 22[214]9[216]-32HB-biotin | ACATAACGCCAACGCACTCCAGCCAG | 5’-biotin |
| 31[98]26[109]-32HB | GAAAGGAACAACTAAACCCATTTTCAGGAGAGGCTATTATTCGAT | |
| 5[116]29[118]-32HB | AACGCGTTAACAAATATCAACTGGAAAGAAAAGTATTAGATAGCA | |
| 15[63]11[65]-32HB | AAGCCTTAGGAATCGCTGTCTTGTTCAGAGA | |
| 2[118]16[112]-32HB | ACCTTTATTAGATAGAAAGGCTCTAAATTCCAACT | |
| 15[126]11[128]-32HB | GCAAGGCATTCAAACATCAATCCCAAAAGTA | |
| 19[109]23[111]-32HB | CCACATGATTGGAAGGTGGAAATTAGCGCGT | |
| 5[95]29[97]-32HB | TTCCGGAATCAATATAACTTATTAGCAATAACTCAAGAGCCACCA | |
| 11[171]8[168]-32HB | ATCACAACCCGTGAGGGGTCAGGCT | |
| 3[28]18[28]-32HB | AATTATCAATAGATTAATTTAAACAATG | |
| 5[53]29[55]-32HB | TGCCGTTATACATGTAAAAAGGTGGCAGATAAGGCGGAGCCACCC | |
| 8[188]5[178]-32HB | GCGCCATGTAAAACGGTACCGAGCTCGATAAC | |
| 11[87]8[84]-32HB | AGAGATAATTAAATTTAAAAATCAT | |
| 16[90]19[87]-32HB | GCGTCTTCGCATTATAATATCAGGA | |
| 25[77]11[79]-32HB | AGAGCCGAAAGCCAAAGGGCGTGTTAGCGCC | |
| 23[112]25[118]-32HB | TTAAAAAGGAAGTCGGTCAGA | |
| 15[105]11[107]-32HB | AATTTTACGCACTCTACCGGATAAGGTAACA | |
| 29[140]15[146]-32HB | CTCAGCATGAGGCTCTGGAAGAATTCTATAGCAAA | |
| 0[214]4[193]-32HB-biotin | TCGAGGTGCCGTAAAGCACTAAATCGCATCACCATCGGCAGGTTTGCCCA | 5’-biotin |
| 2[160]16[154]-32HB | CGAGATAAACCGTCGCAATGCAATAAAGGAAAAGG | |
| 27[182]13[184]-32HB | GTCGAAATACGAAGCAGACCGGATAAGAATT | |
| 4[150]7[153]-32HB | GCGTTAAAGCCTGTTTCCTTGGGTA | |
| 24[41]9[38]-32HB | GAGCCACAATGAAAATCACCACGAG | |
| 26[108]9[101]-32HB | ATTCTGAGGCAATCGGCATTTTCGGAGACTGTATTCATTAAAT | |
| 31[56]26[66]-32HB | TTTTGCTAAACAACTACAACGCGCCACCTTTTGCTGCCCGTAAAGC | |
| 12[94]31[97]-32HB | GTCCGGTATTATGAGCGCGACGGGAAACACTGGAGAATA | |
| 16[132]19[129]-32HB | TAGTAGCCTAAAGTAGCTCAAAGAG | |
| 19[172]23[174]-32HB | CAAATCAAAAAGCGTCCCACTATCGATTCAT | |
| 8[146]5[136]-32HB | ATCGGTGGATTAAGTGTGTGAAATTGTTGGAA | |
| 11[108]8[105]-32HB | AAAGATAAGAAGACCGTGGAGTCAA | |

**Supplementary Table S4.** Modified staples for dye attachment.

| Name | Sequence (5’ -> 3’) | Purpose |
|---|---|---|
| 4[47]2[48]-#9 | GACCAACTAATGCCACTACGAAGGGGGTAGCATTTCACTCCAGGAATATAAAGC | 1 |
| 12[47]10[48]-#6 | TAAATCGGGATTCCCAATTCTGCGATATAATGTTGTTTCTTCGGGCGTACTGAC | 1 |
| 20[47]18[48]-#9 | TTAATGAACTAGAGGATCCCCGGGGGGTAACGTTTCACTCCAGGAATATAAAGC | 1 |
| 4[111]2[112]-#6 | GACCTGCTCTTTGACCCCCAGCGAGGGAGTTATTGTTTCTTCGGGCGTACTGAC | 1 |
| 12[111]10[112]-#9 | TAAATCATATAACCTGTTTAGCTAACCTTTAATTTCACTCCAGGAATATAAAGC | 1 |
| 20[111]18[112]-#6 | CACATTAAAATTGTTATCCGCTCATGCGGGCCTTGTTTCTTCGGGCGTACTGAC | 1 |
| 4[175]2[176]-#9 | CACCAGAAAGGTTGAGGCAGGTCATGAAAGTTTCACTCCAGGAATATAAAGC | 1 |
| 12[175]10[176]-#6 | TTTTATTTAAGCAAATCAGATATTTTTTGTTTGTTTCTTCGGGCGTACTGAC | 1 |
| 20[175]18[176]-#9 | ATTATCATTCAATATAATCCTGACAATTACTTTCACTCCAGGAATATAAAGC | 1 |
| 4[239]2[240]-#6 | GCCTCCCTCAGAATGGAAAGCGCAGTAACAGTTTGTTTCTTCGGGCGTACTGAC | 1 |
| 12[239]10[240]-#9 | CTTATCATTCCCGACTTGCGGGAGCCTAATTTTTTCACTCCAGGAATATAAAGC | 1 |
| 20[239]18[240]-#6 | ATTTTAAAATCAAAATTATTTGCACGGATTCGTTGTTTCTTCGGGCGTACTGAC | 1 |
| 12[111]10[112]-single_mol | TAAATCATATAACCTGTTTAGCTAACCTTTAATTTAAATGCCAAAAAAAAAAAA AAAAGTTTCTTCGGGCGTACTGAC | 2 |
| 9[128]11[127]-single_mol | GCTTCAATCAGGATTAGAGAGTTATTTTCATTTCACTCCAGGAATATAAAGC | 3 |
| 25[77]11[79]-32HB-#6 handle(5') | GTTTCTTCGGGCGTACTGACTTAGAGCCGAAAGCCAAAGGGCGTGTTAGCGCC | 4 |
| 23[49]25[55]-32HB-7nt catching site | TAGCAGCAATCACCCCTCAGATTTTAAATGC | 4 |
| 23[91]25[97]-32HB-7nt catching site | TAGCGTCTCATAGCCATTGACTTTTAAATGC | 4 |
| 4[79]2[80]-7nt-catching-site-up | GCGCAGACAAGAGGCAAAAGAATCCCTCAGTTTTAAATGC | 5 |
| 12[79]10[80]-7nt-catching-site-down | AAATTAAGTTGACCATTAGATACTTTTGCGTTTTAAATGC | 5 |
| 6[79]4[80]-handle#6 | GTTTCTTCGGGCGTACTGACTTTTATACCACCAAATCAACGTAACGAACGAG | 5 |
| 1[160]2[144]-7nt+mismatch-catching-site-left | TTAGGATTGGCTGAGACTCCTCAATAACCGATTTTTAAATGCT | 5 |
| 4[207]2[208]-7nt+mismatch-catching-site-right | CCACCCTCTATTCACAAACAAATACCTGCCTATTTTAAATGCT | 5 |
| 2[175]0[176]-handle#9 | TCACTCCAGGAATATAAAGCTTTATTAAGAAGCGGGGTTTTGCTCGTAGCAT | 5 |
| 23[28]25[34]-32HB-#6 handle | CGTCACCCACCGGACGCCACCTTGTTTCTTCGGGCGTACTGAC | 6 |
| 23[175]25[181]-32HB-#6 handle | CAGTTGATAATCATGCTGCTCTTGTTTCTTCGGGCGTACTGAC | 6 |

Purpose:

(1) 20 nm-grid STORM origami
(2) Single-molecule ATTO647N origami
(3) Single-molecule Cy3b origami
(4) 32HB oscillator origami
(5) 2-color oscillator origami
(6) 32HB axial ruler

Oligos substitute the unmodified oligos with the same name (minus the extension, after the hyphen).

**Supplementary Table S5.** Fluorescent oligos.

| Name | Sequence (5’ -> 3’) | Modification | Purpose |
|---|---|---|---|
| #6-AF647 | CGTCAGTACGCCCGAAGAAAC | 3’ - Alexa Fluor 647 | 1, 6 |
| #9-AF647 | CGCTTTATATTCCTGGAGTGA | 3’ - Alexa Fluor 647 | 1 |
| arm#6-ATTO647N | GTCAGTACGCCCGAAGAAACTTTTTTTTTTTTTTTTGGCATTTA | 3’ - ATTO647N | 2, 4 |
| #9-Cy3b | CGCTTTATATTCCTGGAGTGA | 3’- Cy3b | 3 |
| arm#9-JF549 | GCTTTATATTCCTGGAGTGATTTTTTTTTTTTTTTTGGCATTTA | 3’ Janelia Fluor 549 | 5 |

Purpose:

(1) 20 nm-grid STORM origami
(2) Single-molecule ATTO647N tracking origami
(3) Single-molecule Cy3b tracking origami
(4) 32HB oscillator tracking origami
(5) 2-color oscillator origami
(6) 32HB axial ruler

### Supplementary Note 4: MINFLUX acquisition settings

#### Imaging

| Excitation Scheme | $L_{xy}$ (nm) | $L_z$ (nm) | Photons | Live estimator $\beta_0$ | Live estimator $\beta_1$ |
|---|---|---|---|---|---|
| **3x4 DNA origami** | | | | | |
| 7G | 400 | - | 200 | - | - |
| 4G | 300 | - | 200 | - | - |
| 5D | 150 | - | 120 | 0.90 | 7.18 |
| 5D | 100 | - | 280 | 0.57 | 10.80 |
| 3B | - | 350 | 280 | 0.40 | 2.00 |
| 3B | - | 150 | 280 | 0.40 | 1.50 |
| 3B | - | 100 | 320 | 0.40 | 1.20 |
| DBD | 50 | 75 | 10000 | 0.80 | 22.00 |
| **Nuclear Pore Complex** | | | | | |
| 7G | 400 | - | 200 | - | - |
| 4G | 300 | - | 200 | - | - |
| 5D | 150 | - | 120 | 0.90 | 7.18 |
| 5D | 100 | - | 280 | 0.57 | 10.80 |
| 3B | - | 350 | 400 | 0.40 | 2.00 |
| 3B | - | 150 | 400 | 0.40 | 1.50 |
| DBD | 75 | 75 | 10000 | 0.80 | 11.00 |

#### Tracking

The centering step was conducted in all dimensions using various excitation schemes. The transition between these schemes was determined by a termination condition, which could be either reaching a maximum time ($t_{max}$) or number of photons ($N$), or the live estimated MINFLUX localization being at a specific distance ($d$) from the center of the excitation pattern. In the acquisition of the dual-color origami, two excitation schemes, 640 nm and 560 nm DB, were interleaved in time, with the total acquisition for each wavelength lasting approximately 250 µs.

| | Excitation Scheme | $L_{xy}$ (nm) | $L_z$ (nm) | Termination condition | Live estimator $\beta_0$ | Live estimator $\beta_1$ |
|---|---|---|---|---|---|---|
| | **Axial oscillator** | | | | | |
| Centering | 7G | 400 | - | $t_{max}$: 200 ms<br>$d$: 20 nm | - | - |
| | 5D | 150 | - | $N$: 400<br>$t_{max}$: 100 ms | 0.90 | 7.18 |
| | 3B | - | 350 | $d$: 5 nm | 0.40 | 2.00 |
| Static tracking | DB | 100 | 100 | - | - | - |
| | **Dual-color oscillator** | | | | | |
| Centering (640 nm) | 7G | 400 | - | $t_{max}$: 200 ms<br>$d$: 20 nm | - | - |
| | 5D | 150 | - | $N$: 400<br>$t_{max}$: 100 ms | 0.90 | 7.18 |
| | 3B | - | 350 | $d$: 5 nm | 0.40 | 2.00 |
| Static tracking | DB (640 nm) | 100 | 100 | - | - | - |
| | DB (560 nm) | 100 | 100 | - | - | - |

The scheme codes occurring above are:

| | |
|---|---|
| **7G** | 7 Gaussian at vertices and center of regular hexagon |
| **4G** | 4 Gaussian in a cross |
| **5D** | 4 donuts in a cross plus a central exposure |
| **3B** | 3 bottle beams along the optical axis |
| **DB** | 4 donuts in a cross and 3 bottle beams along the optical axis |
| **DBD**[*] | 4 donuts in a cross plus central donut and 3 bottle beams along the optical axis |

[*]Adding an additional donut exposure in the center allows to make an xy-localization (5 donuts) and a z-localization (3 bottle beams shifted axially) independently.

### Supplementary Note 5: Localization filtering

The parameter $p_0^{xy} = n_0^{xy}/\sum n_i^{xy}$, where $n_i^{xy}$ are the photon counts from the donut exposures in the xy-plane of the excitation pattern with $n_0^{xy}$ being the central donut specifically, gives a measure for how well-centered the last excitation pattern was. The analogue in z is $p_0^z = n_0^z/\sum n_i^z$ depending on the counts from the three bottle beam exposures $n_i^z$. For background we expect $p_0^{xy} = 1/5$ and $p_0^z = 1/3$ since we employ 5 donut beams and 3 bottle beams. The parameter $r_{relative}$ is the distance of the center of the last excitation pattern to the localization made and was used to filter out failed or background localizations that lie outside the excitation pattern.

#### DNA origami

| | |
|---|---|
| $p_0^{xy}$ | [0, 0.1] |
| $p_0^z$ | [0, 0.33] |
| SBR | [0, 10] |
| $r_{relative}$ | [0, 50] nm |
| Brightness | [1.54e4, 4.94e4] counts per second |

#### NPC

| | |
|---|---|
| $p_0^{xy}$ | [0, 0.1] |
| $p_0^z$ | [0, 0.33] |
| SBR | [0, 19.7] |
| $r_{relative}$ | [0, 37.5] nm |
| Brightness | [2e3, 1e5] counts per second |

### Supplementary Figure: Imaging Localization Filtering

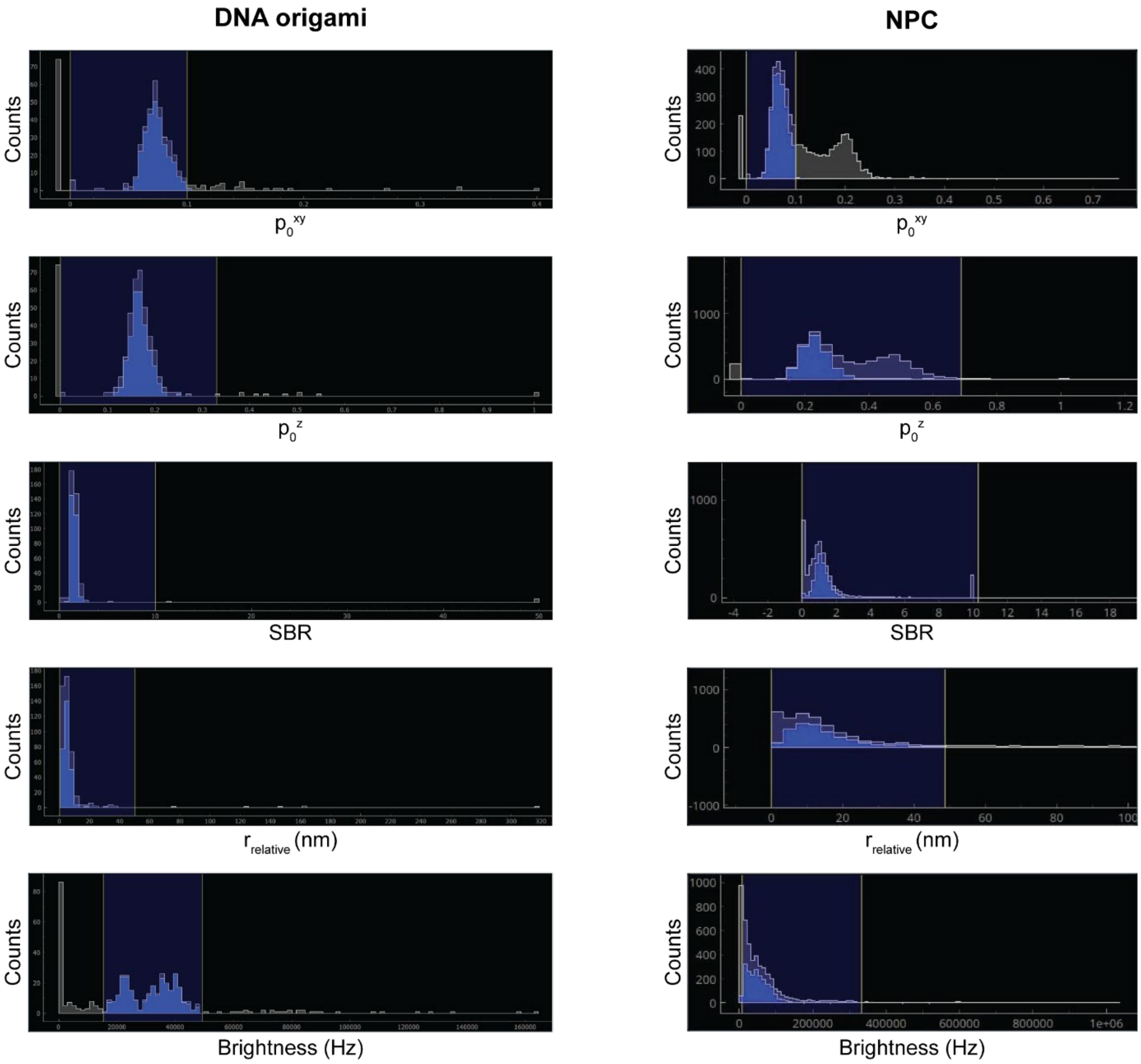

**Supplementary Fig. S17. Imaging data filtering histograms.**

The estimated localizations filtered for data visualization have $p_0$, SBR, $r_{\mathrm{relative}}$ and Brightness in the value ranges shaded in blue.
